# A Physics-Inspired Classical Digital Twin of the Cell: A Composite Multi-Clock Port-Hamiltonian Neural Network Learned from Multi-Omic Circadian Data

**DOI:** 10.64898/2026.07.11.737972

**Authors:** Dibakar Sigdel, Namuna Panday

## Abstract

We present a physics-inspired classical digital twin of the cell: a graph neural network constrained to a compartmental, multi-clock port-Hamiltonian form, with parameters learned from multi-omic measurements. The port-Hamiltonian structure is a modelling choice — it buys conservation, passivity and a clean separation of storage, routing and dissipation — not a claim about what a cell is. The state pairs each species’ abundance deviation with a phase coordinate, assigned only where a per-clock rhythmicity gate certifies periodicity. Stored energy decomposes over five functional compartments, so stability is verified compartment by compartment. Two distinct clocks are included — the 24-hour transcription–translation loop and the 20-hour transcription-independent redox oscillator — coupled through a zero-net-power link, with the central-dogma correspondence hard-wired and moiety pools exact invariants. On a real mouse-liver three-omic dataset the verdict is mixed. Across ten seeds the trained twin is thermodynamically stable (no violations at any sampled state) and forecasts held-out segments (root-mean-square error 0.325 ± 0.002). Its central prediction — cross-omic phase lag equals arctan of clock frequency over degradation rate — matches the aggregate transcript-to-protein lag (5.74 ± 0.03 versus 4.90 hours), but the per-gene correlation is indistinguishable from zero (*r* = 0.06 ± 0.27, sign unstable across seeds), so the law is supported in aggregate and unresolved per species. Recovery of withheld interaction edges is at chance (AUROC 0.50 ± 0.13, nine of ten seeds scoreable): 24 timepoints do not identify network topology, which we report as a bound on what this data volume supports rather than as a property of the framework. Because the port-Hamiltonian form is imposed by construction, edits to the twin preserve it, so specialisation and disease can be expressed as structured perturbations of this reference twin rather than as separate models.

## I. INTRODUCTION

Modern systems biology aspires to derive mechanistic, predictive models of the living cell from multi-omic data, yet faces a fundamental tension: the technologies that measure genomics, transcriptomics, proteomics, and metabolomics produce static, high-dimensional snapshots, while the cell is an open, thermodynamically driven dynamical system evolving in continuous time. Bridging that gap — turning molecular inventories into a governing dynamics — is the central problem this work addresses.

### A. From Whole-Cell Models to the Virtual Cell

The most complete mechanistic account of a cell to date is the whole-cell model of *Mycoplasma genitalium*, which reproduced observable phenotype from genotype by integrating twenty-eight functional submodels of distinct cellular processes [1]. Its successors extended this program to *Escherichia coli* through mechanistic simulation cross-evaluated against heterogeneous datasets [2] and to a genome-scale kinetic model of a minimal cell whose emergent behaviours arise from first-principles biochemistry [3]. These efforts established both the promise and the cost of hand-built mechanistic modelling: each submodel is curated individually, parameterised from disparate experiments, and stitched together by expert judgement [4]. The approach does not scale to the mammalian cell, and it does not learn from data.

The contemporary response is the *AI virtual cell* : a data-driven, learnable surrogate for cellular state and behaviour [5, 6]. Single-cell foundation models — transformer architectures pre-trained on tens of millions of transcriptomes such as scGPT [7] and Geneformer [8] — have demonstrated transferable representations of cell state and *in silico* perturbation response. Yet these models are fundamentally *static and correlational* : they embed snapshots and predict masked features, but they carry no explicit time axis, no conservation law, and no thermodynamic constraint. A learned representation that violates mass balance or manufactures free energy can still score well on a reconstruction benchmark while being physically impossible as a dynamics.

### B. Physics-Informed and Structure-Preserving Learning

A parallel line of work embeds physical law directly into the learning problem. Physics-informed neural networks penalise violation of a governing equation [9, 10], while *structure-preserving* architectures build conservation and geometry into the model class itself: Hamiltonian and Lagrangian neural networks learn dynamics that conserve a learned energy by construction [11, 12], and Hamiltonian graph networks extend this to many-body systems on a graph [13]. In systems biology, thermodynamic constraints have long been imposed at steady state through constraint-based methods — flux balance analysis and its energy-balance extension rule out thermodynamically infeasible flux distributions [14, 15] — and geometric approaches recast cell-fate decisions as motion on a low-dimensional landscape or vector field [16, 17]. What has been missing is a single framework that is at once *learnable from multi-omic data*, *dynamical in continuous time*, and *thermodynamically honest by construc- tion* — combining the data-driven expressiveness of the virtual-cell program with the built-in physical guarantees of structure-preserving learning.

### C. This Work

We close that gap with a *port-Hamiltonian* formulation of cellular dynamics learned by a graph-neural-network surrogate. Most deep-learning approaches to gene-regulatory-network (GRN) inference or trajectory modelling treat multi-omic data as unstructured feature vectors, ignoring the physical laws — mass-action kinetics, conservation of matter, irreversibility of dissipation — that constrain real cellular processes. A biologically coherent framework instead requires three ingredients simultaneously: (i) a *geometry* that reflects the oscillatory, phase-locked nature of biological programs; (ii) a *dynamics* that enforces thermodynamic structure, cleanly separating conservative regulatory routing from irreversible molecular decay; and (iii) a *surrogate* expressive enough to capture the high-dimensional, non-linear, state-dependent rewiring of the GRN under perturbation. The port-Hamiltonian framework supplies (ii) by construction, the phasor/torus geometry supplies (i), and the GNN supplies (iii).

### D. Biological Clocks and Phasor Representation

Cells are not static chemical reactors; they are *dissipative oscillators*, and they run more than one clock. The circadian clock (*T* ≈ 24 h) is a transcription–translation feedback loop [18]; the redox clock is a transcription-*independent* peroxiredoxin oxidation rhythm that persists even in enucleated cells [19, 20]. These are mechanistically distinct oscillators coupled through metabolism (the NAD^+^/SIRT1 axis [21]), not two harmonics of one clock. Near a Hopf bifurcation an oscillatory pool admits a universal reduced description — the Stuart–Landau normal form **[22]:**

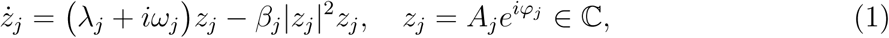

where *λ_j_ >* 0 is the growth rate, *ω_j_* the natural frequency drawn from the appropriate clock, and *β_j_*|*z_j_*| *z_j_* the amplitude-saturating term that fixes the stable limit cycle at 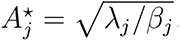.

The phasor *z_j_* is a *derived* readout of a genuinely oscillatory pool (Section II), and the rhythmic pools of each clock live on that clock’s torus.

### E. Port-Hamiltonian Dynamics for Cell Biology

Port-Hamiltonian (pH) systems [23] provide the mathematical language in which energy storage, dissipation, and exchange with the environment are separated by construction (Fig. 1 sets out the whole construction at a glance):

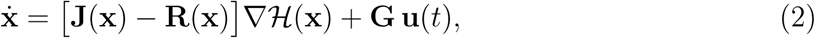

**FIG. 1:**
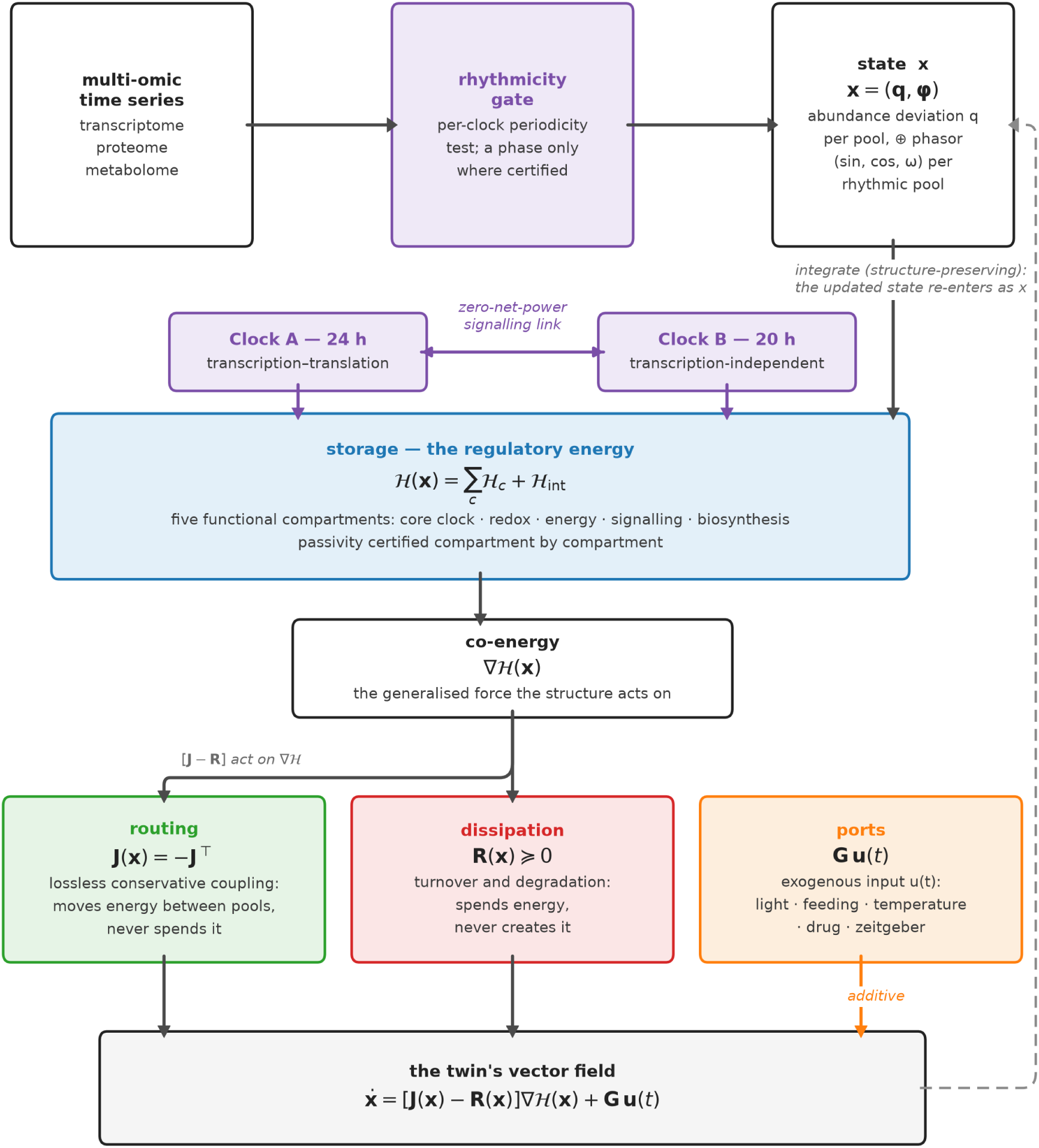
The classical port-Hamiltonian digital twin of the cell. A per-clock rhythmicity gate decides which pools earn a phase coordinate, so **x** pairs an abundance deviation with a phasor only where periodicity is certified. Storage H (blue) decomposes over five compartments driven by two clocks (purple) coupled through a zero-net-power link; its gradient is the force on which **J** = −**J**^⊤^ (green) and **R** ≥ 0 (red) act, while **Gu**(*t*) (orange) adds exogenous drive. The central-dogma map, conserved-moiety pools, and the structure of **J** and **R** are imposed, not penalised. See Sec. II.

where H(**x**) is the regulatory energy (Lyapunov storage function), **J** = −**J**^⊤^ encodes lossless conservative coupling (reversible biochemical reactions, complex formation), **R** = **R**^⊤^ ≥ 0 encodes irreversible dissipation (mRNA degradation, protein turnover, metabolic heat loss), and **G u**(*t*) represents exogenous ports (Zeitgeber signals, drug dosing, nutrient influx). The resulting power balance Ḣ = −∇H^⊤^**R**∇H + **y**^⊤^**u** ≤ **y**^⊤^**u**, written throughout in terms of the conjugate port output **y** = **G**^⊤^∇H, is the mathematical statement of *passivity* : the cell cannot generate free energy from its regulatory network alone.

### F. Limitation of Static Connectomes

A critical but underappreciated failure mode of prior pHNN formulations is the use of *static* interaction matrices. A living cell’s regulatory network is state-dependent: the same molecular network routes couplings differently under pharmacological perturbation than at homeostasis. A static **J** captures only the time-averaged connectome, missing the non-linear, context-specific regulatory logic that governs cellular resilience. We therefore retain the state-dependent connectome **J**(**x**) predicted by the GNN surrogate, now assembled block-wise over functional compartments.

### G. Contribution

We present the **Cellular GNN-pHNN**, a framework that addresses these limitations through a composite, multi-clock, compartmental port-Hamiltonian model learned by a GNN surrogate. Our contributions are:

1. A **two-layer state** in which abundance deviation *q_j_* (directly measured) is primitive and the phasor *z_j_* is derived only for gated rhythmic pools, replacing the biologically false assumption that every pool is an inertial oscillator.
2. A **composite compartmental Hamiltonian** H = ∑*_c_* H*_c_* +H_int_ whose compartments are functional biological modules (core clock, redox, energy, signalling, biosynthesis), each carrying its own clock, so the model’s energy decomposition maps onto biological processes rather than measurement layers.
3. A **clock bank** of mechanistically distinct oscillators (transcription-dependent circadian TTFL and transcription-independent redox rhythm) coupled through a zero-net-power NAD^+^/SIRT1 signalling port.
4. A **per-clock rhythmicity gate** that assigns each pool to a clock from its spectrum and estimates a time-referenced acrophase, making clock membership an inference from data.
5. A **structure-preserving connectome and dissipation**: block-diagonal intra- compartment **J***_c_* plus inter-compartment mass bonds and modulated ports, with a diagonal positive dissipation **R** = diag(*r_j_*) ≥ 0 initialised from measured degradation rates.
6. A **composite physics loss** with global *and* per-compartment passivity, and a PLV term demoted from a fitting target to a weak prior, so that recovery of coupling structure is a measured outcome rather than an injected one. Measured on this dataset, that recovery does not occur: edge recovery is at chance, which we report as a bound on what 24 timepoints identify (Section IV).
7. A **doubly-falsifiable phase-cascade prediction**: the circadian transcript→protein lag and the redox protein→metabolite lag are predicted from the *same* learned dissipation operator via tan Δ*φ* = *ω/k*_deg_, with each clock’s frequency. Tested here, the first arm is confirmed in aggregate and unresolved per gene; the second is not scoreable on the available metabolome sampling.
8. A **reference twin that admits structured perturbation**: because the port-Hamiltonian form is imposed by parameterisation rather than by penalty, edits to topology, dissipation, set-point or port structure leave passivity intact, so specialisation and disease can be built on this twin as class-preserving deformations of it rather than as separate models (Section II H). Developing the edit algebra, and applying it to specialisation and to disease courses, is left to future work.

*a. Positioning.* The framework occupies ground that none of the three established camps reaches alone. Unlike hand-built whole-cell models [1, 2], it is learned end-to-end from data and does not require a curated mechanism for every process. Unlike single-cell foundation models [7, 8], which embed static snapshots, it is a continuous-time *dynamics* with an explicit energy, dissipation, and input port — so passivity, mass conservation, and moiety invariants hold by construction rather than being hoped for. And unlike generic structure-preserving networks [12, 13], whose conserved quantity is an abstract learned scalar, its Hamiltonian is decomposed over named biological compartments and its dissipation operator is tied to measured molecular turnover, yielding a prediction — the cross-omic phase cascade — that is quantitative, mechanistic, and falsifiable against experiment. The result is, to our knowledge, the first virtual-cell surrogate that is simultaneously data-driven, dynamical, and thermodynamically certified. Certified here means the thermodynamic properties hold by construction and are verified numerically; it does not mean the twin is quantitatively predictive at species resolution, which on this dataset it is not.

*b. What this model is, and what it is not.* The object developed here is a *physics-inspired classical digital twin* of the cell: a neural network whose architecture is constrained to the port-Hamiltonian form and whose parameters are learned from multi-omic measurements. The port-Hamiltonian structure is a modelling choice, adopted because it supplies conservation, passivity and a clean separation of storage, routing and dissipation — properties that make a learned model interrogable and its failures diagnosable. It is not a claim about what a cell is. Living cells are open, stochastic, spatially organised and almost certainly not exactly Hamiltonian at any level of description; the value of the present construction is that it *mimics* measured cellular behaviour well enough to be probed under conditions the experiment did not visit, while carrying physical guarantees by construction rather than by penalty. Every result below should be read as a statement about the twin, and about how faithfully the twin tracks the data — never as a statement that biology obeys these equations.

*c. A note on the word “cell”.* Throughout this paper, unless the context is explicitly biological, *cell* means the classical port-Hamiltonian digital twin of the biological cell — the learned model, not the living object. This convention lets the exposition read naturally without repeating the qualifier: statements about “the cell’s” energy, connectome or set-point are statements about the twin. Where a sentence concerns the biological cell itself, it says so.

## II. THEORETICAL FRAMEWORK

### A. Two-Layer State: Abundance and Derived Phasor

Let the cell contain *N* molecular pools indexed *j* = 1*, …, N* spanning three omic layers: Genomics/Transcriptome (*j* ∈ G, |G| = 40), Proteome (*j* ∈ P, |P| = 35), and Metabolome (*j* ∈ M, |M| = 25). The primitive state variable is the **abundance deviation** *q_j_* of each pool from its homeostatic set-point — the quantity that omics technologies actually measure. Its conjugate variable is a chemical potential *µ_j_* = *∂*H*/∂q_j_*, so that (*q_j_, µ_j_*) form a genuine power-conjugate pair with a conserved, measurable base variable. This avoids a mechanical (*φ_j_, ω_j_*) phase–frequency pairing, which would treat each pool as if it carried inertia — a molecular abundance has no momentum.

The phasor coordinate is *derived*, not primitive. For a pool that genuinely oscillates, we write

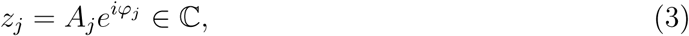

a reduced readout of the oscillatory subspace obtained *after* a rhythmicity gate (Section III) certifies that the pool oscillates. Non-rhythmic pools carry no phase coordinate; they participate in the dynamics through their abundance alone. Only the *N_r_* rhythmic pools are placed on the torus, so the phase manifold is T*^Nr^*, not T*^N^* .

### B. A Composite, Multi-Clock, Multi-Compartment Hamiltonian

The cell is not organised by measurement layer but by *function*. We decompose the state into *C* functional **compartments**, each a group of molecular pools that participate in one coherent biological process. Compartment *c* carries its own storage function H*_c_*, and the total storage is a composite Hamiltonian

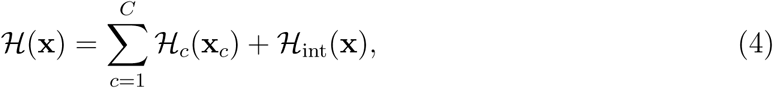

where **x***_c_* is the state restricted to compartment *c* and H_int_ collects genuine inter-compartment couplings. This is the port-Hamiltonian expression of a modular cell: each compartment is a sub-system with its own internal energy, interconnected through ports.

Crucially, the compartments do not all march to the same clock. We equip the model with a **clock bank** — a small set of biologically distinct oscillators, each with its own angular frequency:

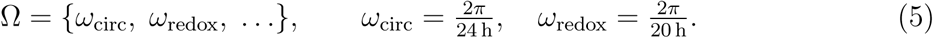

The two clocks are genuinely distinct in mechanism, not merely in period. The *circadian* clock is the transcription–translation feedback loop (TTFL), an autoregulatory gene circuit [18]. The *redox* clock is the transcription-*independent* peroxiredoxin oxidation rhythm, which persists in enucleated cells and even in the absence of transcription [19, 20]. A rhythmic pool belongs to exactly one clock; its phase lives on that clock’s circle, so the joint phase manifold factorises as a product of per-clock tori 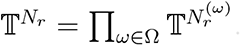

### C. Rhythmicity Gating and Clock Assignment

Clock phase is treated as both a model state and an observation coordinate, but *only* for genuinely oscillatory features. A per-clock rhythmicity gate (Section III) tests each pool against each clock’s spectral band, assigns it to the clock whose band contains its dominant period, and estimates a *time-referenced* acrophase by least-squares fit at that clock’s frequency. Pools that fail every band remain pure abundance variables. This makes clock membership an inferred property of the data, not an imposed label.

### D. Two Structurally Distinct Edge Types

The interconnection structure distinguishes two mathematically distinct kinds of edge, following compositional port-Hamiltonian theory [23]:

*a. Mass bonds* are power-conjugate couplings that carry real flux (transcription →translation, enzymatic conversion). They enter the interconnection matrix **J** skew- symmetrically and are therefore exactly power-conserving: ∇H^⊤^**J** ∇H = 0 for any **J** = −**J**^⊤^. The central-dogma gene→protein correspondence is hard-wired as a near-diagonal mass-bond block.

*b. Modulated ports* are state-dependent gains that transmit *information*, not mass (allosteric regulation, signalling). They are implemented as skew-symmetrised modulated terms Γ(**x**) − Γ(**x**)^⊤^ so that their net power contribution is identically zero, yet the gain Γ depends on the state. The circadian↔redox coupling (mediated biologically by the NAD^+^/SIRT1 axis that links metabolic redox state to the core clock [21]) is exactly such a zero-net-power modulated port — a signalling link, not a conservative mass bond.

### E. Stoichiometric Conservation

A stoichiometric matrix **S** encodes moiety conservation exactly: the adenylate pool (ATP+ADP+AMP) and the redox cofactor pools (NAD^+^/NADH) are sum-locked invariants of the learned dynamics, **Sq̇** = 0. A conserved pool oscillates only as a *ratio*, never as independent abundances; we therefore place the conserved moiety pools in a clockless, homeostatically buffered compartment, and reserve the rhythmic cascade readouts for free (non-conserved) metabolites. This prevents the conservation constraint and the phase-cascade prediction from contradicting each other.

### F. Composite Port-Hamiltonian Dynamics

The complete dynamics retain the port-Hamiltonian form of Eq. (2), with the interconnection and dissipation operators assembled block-wise from the compartments:

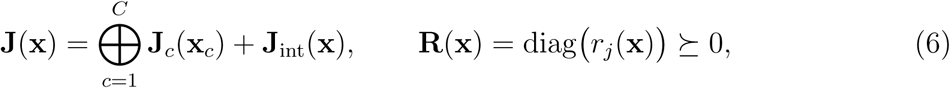

where ⊕ is the block-diagonal (intra-compartment) sum, **J**_int_ carries the inter-compartment mass bonds and the skew-symmetrised clock-coupling ports, and the per-pool dissipation rates *r_j_*(**x**) = softplus(·) *>* 0 make **R** ≥ 0 at every state. The dissipation acts on the abundance block and is initialised from measured degradation rates *k*_deg_: dissipation *is* molecular turnover.

### G. Thermodynamic Power Balance and Per-Compartment Passivity

The central invariant is the power balance

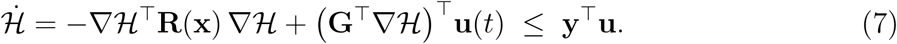

Without external input, Ḣ ≤ 0 by **R** ≥ 0: the network can only dissipate stored “energy” (a Lyapunov storage function, not joules). Because the storage is composite (Eq. 4), passivity can be certified compartment by compartment: restricting ∇H and **R** to a compartment’s abundance block, each local dissipation 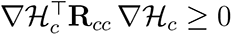 must hold. The skew-symmetrised clock-coupling ports contribute exactly zero to Ḣ, so inter-clock signalling never manufactures free energy. This per-compartment passivity is a strictly tighter thermodynamic check than the global one, and it is enforced directly in the training loss.

### H. Structured Perturbations: Specialisation and Disease

The reference twin is a single object, and the biology it must eventually cover is a family. Both cell specialisation and disease are naturally expressed in this framework as *class-preserving* deformations of the reference quadruple (**J**, **R**, **Q**, **B**): edits that change topology, dissipation, set-point or port structure while leaving the port-Hamiltonian form — and therefore passivity — intact by construction. A specialised cell is then a different fixed point reached by a legal edit rather than a different physics, and a disease is a trajectory through the same edit space.

That programme is not developed here, and we state the reasons for the boundary rather than merely drawing it. Specialisation and disease are questions about a *family* of twins: which edits are admissible, what the composition of two edits reaches, and which lesions are reversible through existing ports. Each requires an edit algebra and an account of its path-ordering — the observation that composing two edits can depend on the order in which they are applied — neither of which the present data constrain. The scope of this paper is the reference twin itself: its construction, and what a real multi-omic time course does and does not identify about it.

## III. METHODS

### A. Functional Compartments and the Clock Bank

The *N* = 100 molecular pools are partitioned into five *functional compartments*, each a biological process rather than a measurement layer (Table I). Two compartments carry clocks: the core-clock compartment runs on the circadian oscillator, the redox compartment on the redox oscillator; the energy, signalling, and biosynthesis compartments are clockless. Every omic layer contributes rhythmic pools (genomics and proteome to the circadian clock, proteome and metabolome to the redox clock), so no clock is confined to one measurement layer.

**TABLE I:** The five functional compartments, their clocks, member pools, and biological roles. The three conserved moiety pools (adenylate, NAD, cofactor) are placed in the clockless energy compartment; the redox clock drives free, non-conserved metabolites so its cascade lag is cleanly measurable.

| Compartment | Clock | Members | Role |
| --- | --- | --- | --- |
| core_clock | circadian | $\mathcal{G}[0:12], \mathcal{P}[0:12]$ | TTFL gene circuit |
| redox | redox | $\mathcal{P}[12:20], \mathcal{M}[12:20]$ | redox enzymes + free metabolites |
| energy | none | $\mathcal{P}[20:28], \mathcal{M}[0:12]$ | conserved moiety pools |
| signalling | none | $\mathcal{G}[12:24], \mathcal{P}[28:35]$ | signal transduction |
| biosynthesis | none | $\mathcal{G}[24:40], \mathcal{M}[20:25]$ | anabolic pathways |

### B. The Emergent Phase-Cascade Prediction

The model makes a falsifiable, model-independent prediction about cross-omic timing. When an upstream rhythmic pool (e.g. a transcript) drives a downstream pool (its protein) that relaxes as a first-order system with degradation rate *k*_deg_, the downstream oscillation lags the upstream one by a phase set entirely by the turnover rate and the clock frequency,

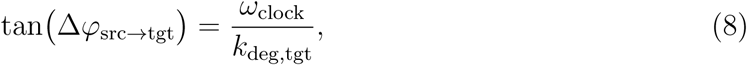

with *no* lag inserted by hand. Because *ω*_clock_ is fixed by the biology and *k*_deg_ can be taken from independently published degradation half-lives, Eq. 8 predicts the measured transcript →protein phase lag without fitting any parameter to the lag itself — the basis of the model-independent test in Section IV. Throughout, conserved moiety pools are renormalised at each timestep to enforce **Sq** = const exactly.

### C. Real Multi-Omic Data

The model is fit to and evaluated on an assembled tri-omic mouse-liver circadian dataset built entirely from public repositories. The three layers are matched by gene/protein symbol and aligned on circadian phase (CT/ZT), not same-animal sampling:

- **Transcriptome:** Zhang et al. 2014, “A circadian gene expression atlas in mammals” [24], deposited at NCBI GEO under accession GSE54650 [25] (Affymetrix Mouse Gene 1.0 ST); liver, CT18–64 at 2 h resolution (two circadian cycles, 22,105 genes after probe-to-gene collapse). Clock-gene amplitudes confirm strong circadian structure (Nr1d1 27.8×, Dbp 20.5×, Arntl 11.9× fold).
- **Proteome:** Robles, Cox & Mann 2014 [26], PLoS Genetics 10(1):e1004047, SILAC mouse liver; CT0–45 at 3 h resolution, 3,072 proteins. This dataset also supplies author-measured transcript→protein phase lags (Table S4, 139 matched genes), used below as a *model-independent* cascade target.
- **Metabolome:** mouse-liver metabolome time-series deposited by Meng et al. at the NIH Metabolomics Workbench under accession ST002079 [27] (12-hour-clock/lipid-metabolism study), ZT0–20 at 4 h resolution (one 24 h cycle, 6 timepoints), 3,552 metabolite/lipid features.

Raw layers arrive on incomparable scales (array intensity, log_2_ SILAC ratios, LC–MS intensity) and are placed on a common *O*(1–10) abundance scale by a monotone per-layer transform (log_2_ for intensity layers, then a robust affine to a shared median/IQR), which preserves each pool’s phase and relative amplitude. Each layer is fit with a single-harmonic cosinor at the 24 h period, and the model’s fixed 40/35/25 compartment slots are populated by real pools: core-clock slots by canonical clock genes, cascade slots by the Robles-matched transcript→protein pairs (27 co-selected pairs), redox slots by rhythmic redox enzymes, and the remainder by the most rhythmic pools per layer. Node-wise degradation rates *k*_deg_ are drawn from published half-lives [28] (*k* = ln 2*/t*_1_*_/_*_2_; mRNA median *t*_1*/*2_ ≈ 7 h, protein ≈ 46 h), used as order-of-magnitude layer priors rather than per-gene measurements. Each layer derives from the public accessions cited above, from which its full retrieval provenance can be traced.

*a. Cross-cohort and sampling caveats.* The three layers come from different cohorts and platforms; only same-phase, not same-animal, correspondence is assumed. The metabolome timecourse (6 points over a single 24 h cycle at 4 h spacing) cannot resolve a 24 h period against nearby periods, which limits the internal redox cascade test (Section IV).

### D. Per-Clock Rhythmicity Gate

The rhythmicity gate tests each pool against every clock (Fig. 7). For each clock *ω* it applies a Butterworth bandpass over a *non-overlapping* detection band (circadian 22–26.5 h, redox 18–22 h, split at ∼22 h) and scores dominant periodicity by Lomb–Scargle periodogram [29, 30] over the union band. A pool passing the SNR and amplitude thresholds is assigned to the clock whose band contains its dominant period, and its acrophase is estimated by a *time-referenced* least-squares fit,

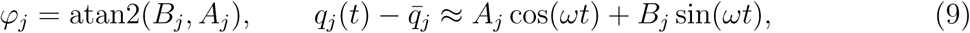

which preserves the *t* = 0 phase reference the cascade test requires. Pools that fail every band remain non-rhythmic abundance variables.

### E. Network Architecture

Figure 2 shows the whole construction. Four networks are trained jointly, and the division of labour between them is the design: one learns a scalar energy, three learn the operators that act on its gradient.

**FIG. 2:**
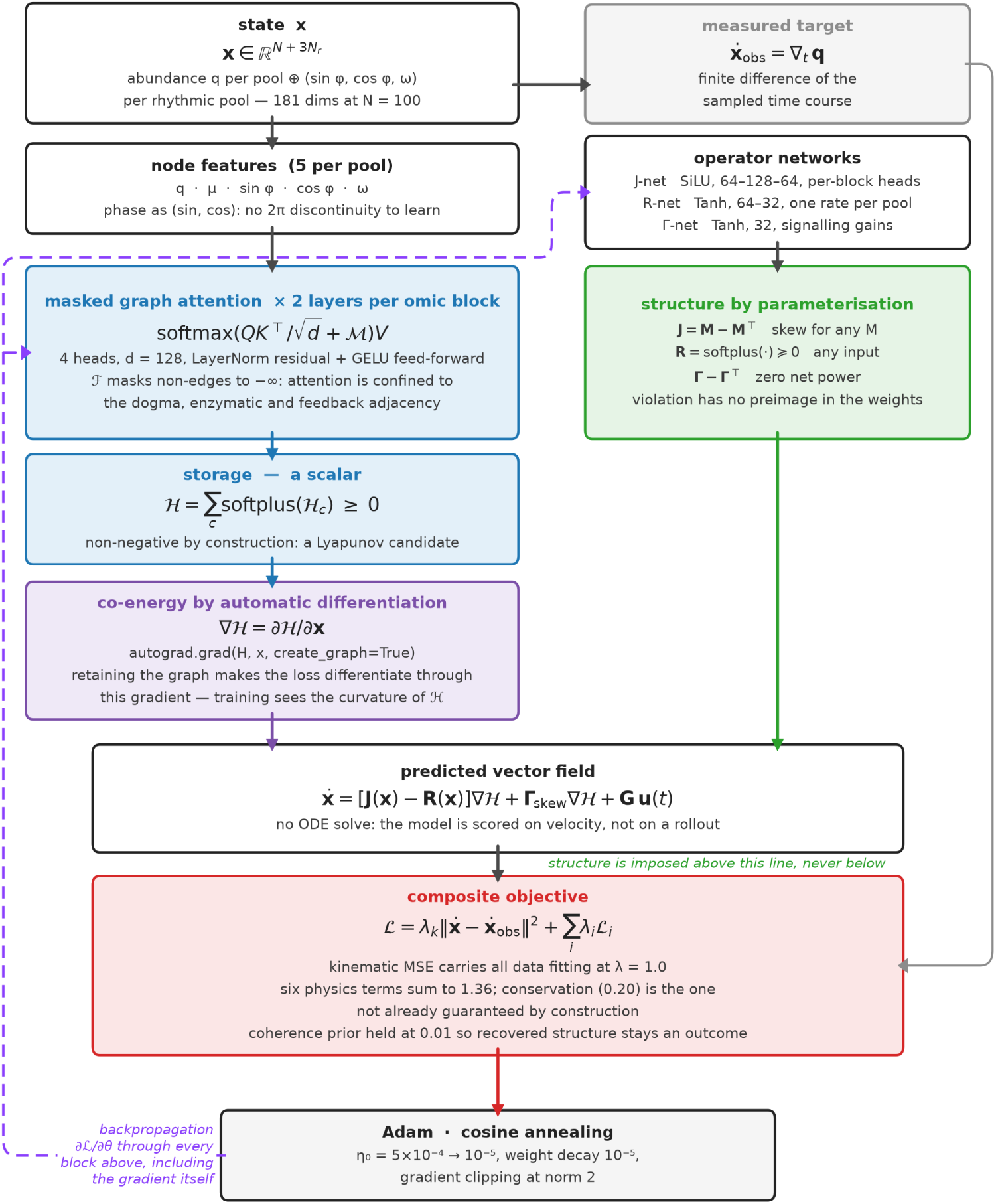
Architecture, forward path and reverse path. Four networks are trained jointly and only one is scalar-valued. *Left:* the state is embedded per pool and passed through masked graph attention to a single non-negative number, the storage H; the predicted velocity is the *gradient* of that number. *Right:* three smaller networks emit the operators, each through a construction that makes its physical property unconditional rather than penalised. *Purple, dashed:* the reverse path. Because the co-energy block is itself differentiable (`create graph=True`), the loss backpropagates *through* the gradient, so training constrains the curvature of H and not only its slope. The measured derivative enters at the objective alone: it is a target, never an input.

*a. The energy network.* The storage function is the composite Hamiltonian of Eq. (4). Each pool enters as a five-dimensional node feature—abundance deviation *q_j_*, chemical potential *µ_j_*, and, for pools the gate certifies as rhythmic, the triple (sin *φ_j_,* cos *φ_j_, ω_j_*).

Phase is carried as its sine and cosine rather than as an angle so that the wrap at 2*π* is not a discontinuity the network must learn around. Two graph-attention layers per omic block follow, with four heads at *d_h_* = 128, LayerNorm residual connections and a GELU feed-forward block, computing

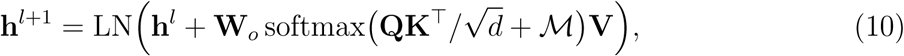

where M sets the score to −∞ on every non-edge of the biological adjacency. Attention is therefore confined to the central-dogma, enzymatic and feedback blocks: a transcript cannot attend to an unrelated metabolite whatever the data suggest. This is the same commitment as the skew construction below—a constraint carried by the architecture rather than by a regularisation term. Layer sub-energies pass through a softplus before summation, so H = ∑*_c_* H*_c_* ≥ 0 is a Lyapunov candidate by construction. A per-compartment readout head pools node embeddings by compartment mask to yield the individual H*_c_* as an interpretable diagnostic; the dynamics use the total.

*b. The operator networks.* Three smaller networks read the same state. **J** is produced by a SiLU trunk (64−128−64) with one head per block, emitting only upper-triangle entries; **R** by a Tanh network (64 − 32) emitting one rate per pool; and the modulated gains **Γ** by a Tanh network of width 32. None of the three is free to violate its physical property, because the property is imposed on the output by construction: **J** = **M** − **M**^⊤^ is skew for any **M**, **R** = softplus(·) is positive for any input, and **Γ** − **Γ** transmits exactly zero net power. A violating configuration has no preimage in the weights, so the guarantee holds at initialisation, at every step, and for inputs far outside the training distribution—which a penalty term cannot promise.

*c. Forward path.* The network never emits a velocity. It emits the scalar H, and the velocity is obtained by differentiating it, ∇H = *∂*H*/∂***x**, before the operators act:

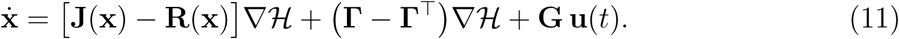

Every prediction is thus a gradient field of a non-negative scalar, which is what makes “conservative routing” a property of the model rather than a description of it.

*d. Reverse path.* Training is ordinary reverse-mode automatic differentiation with Adam; the structure-preserving content lies entirely in the forward construction. One detail is not ordinary. The co-energy is computed with the gradient graph retained, so the loss differentiates *through* ∇H—a second derivative of the storage function. Training therefore constrains the curvature of H, not only its slope, which is what renders the energy landscape identifiable rather than merely the velocity field it induces.

*e. What is not in the loop.* No ODE solve occurs during training. Each step draws a batch of time points, not trajectories, and scores the predicted velocity against a finite difference of the measured series. The consequence is stated plainly because it bounds the claims: the model is fitted to be locally correct about ẋ, and every trajectory-level statement is an extrapolation from that. Section VI A returns to the point.

### F. Compartmental Connectome and Dissipation

The interconnection operator is assembled block-diagonally over compartments plus an inter-compartment block (Eq. 6). Intra-compartment skew blocks are masked to the sparse biological adjacency; the central-dogma gene→protein mass bond is hard-wired near-diagonal. The circadian↔redox clock coupling enters as a strictly upper-triangular modulated gain Γ(**x**) that is skew-symmetrised at use, Γ − Γ^⊤^, contributing exactly zero net power. Dissipation acts on the abundance block as a diagonal, strictly positive rate matrix

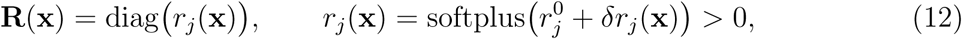

so **R** ≥ 0 holds trivially from the positive softplus entries. The base rates 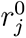 are initialised from measured degradation rates (via the inverse softplus of *k*_deg_*_,j_*), so **R** starts at biological turnover, and the state-dependent correction *δr_j_*(**x**) modulates the effective rate while preserving positivity.

### G. Composite Physics Loss

Training minimises a weighted composite loss

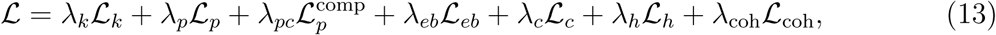

with L*_k_* the kinematic MSE, L*_p_* the global passivity penalty E[ReLU(Ḣ)], and 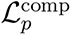 a *per-compartment* passivity penalty

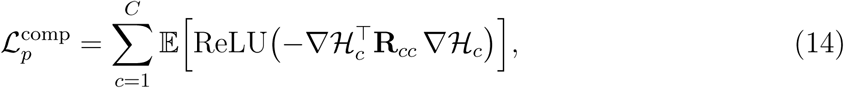

which enforces that *each* compartment dissipates — a tighter check than the global one. L*_eb_* closes the power balance (Eq. 7), L*_c_* enforces stoichiometric conservation ‖**Sq̇**^2^‖, L*_h_* pins the homeostatic set-point, and

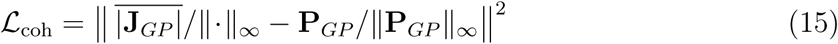

is the phase-locking-value (PLV) coherence term. Its weight is deliberately kept small, *λ*_coh_ = 0.01, so that it acts as a *weak prior* rather than a fitting target: the learned coupling is not driven to the PLV matrix, and any recovery of coherent structure is a measured outcome, not an imposed one (Section IV). Default weights are (*λ_k_, λ_p_, λ_pc_, λ_eb_, λ_c_, λ_h_, λ*_coh_) = (1.0, 0.5, 0.25, 0.3, 0.2, 0.1, 0.01). Optimisation uses Adam with cosine annealing (*η*_0_ = 5×10*^−^*^4^, *η*_min_ = 10*^−^*^5^) and gradient clipping (max norm 2).

### H. State Dimensions, Parameter Budget, and Computational Complexity

It is useful to state plainly the sizes of the objects the model carries, both to make the method reproducible and to locate precisely where a full whole-cell model would become expensive. The model works in *vector and matrix* form throughout: no higher-order tensor state is introduced.

*a. State and operator dimensions.* Each of the *N* molecular pools carries an abundance deviation *q_j_* and its conjugate potential (with a phasor coordinate *derived* for the rhythmic subset after gating, not added as an extra stored axis), so the dynamical state is a real vector **x** ∈ R*^N^*^+3*N*^*^r^* : one abundance coordinate per pool, and three phasor coordinates (sin *φ_j_,* cos *φ_j_, ω_j_*) for each of the *N_r_* pools that pass the rhythmicity gate. The conjugate potential *µ_j_* = *∂*H*/∂q_j_* is *derived* from the storage function rather than carried as an independent coordinate, so it does not enter the state count. For the *N* =100 mouse-liver network (|G|=40, |P|=35, |M|=25) the gate returns *N_r_* = 27 rhythmic pools (Section IV A), so **x** ∈ R^181^ with 3*N_r_* = 81 phasor coordinates. The interconnection and dissipation operators act on this state; **J** is skew and masked to the sparse biological adjacency (only O(*N* + |*E*|) nonzeros over the |*E*|≈450 central-dogma and regulatory edges (445–459 over the seeds sampled), while **R** = diag(*r_j_*) is diagonal with O(*N*) nonzeros. The storage function H : ℝ*^N^*^+3*N*^*^r^* → ℝ is a scalar; its gradient ∇H is the field the dynamics integrate. The exported operator matrices are written in the (*q, µ*) conjugate-pair convention on the abundance block, giving 200 × 200 arrays whose dissipative content occupies the 100 potential indices; the network’s internal state is the 181-dimensional vector above.
*b. Parameter budget.* The trainable parameters are dominated not by the operators but by the GNN energy surrogate that produces H: with *d_h_*=128 hidden dimensions, the attention-based node and edge embeddings account for the great majority of the roughly 1.6 × 10^6^ parameters of the *N* =100 model. The physically-structured pieces are comparatively tiny — the connectome mask, the diagonal dissipation rates initialised from measured degradation, and the clock-coupling gain together scale only as O(*N* + |*E*|). The parameter count is therefore *not* quadratic in *N* : it is a fixed-size surrogate core plus a term linear in the biological graph. (The O(*N* ^2^) figure that does appear below is the *dense-operator storage*, a memory cost, not a trainable-parameter count — the two should not be conflated.)
*c. Computational complexity.* Assembling and storing the dense operators is O(*N* ^2^) in memory (4 × 10^4^ entries at *N* =100), and each forward step — evaluating ∇H, forming (**J** − **R**)∇H, and integrating — is O(*N* ^2^) per step in the dense implementation (reducible to O(*N* + |*E*|) by exploiting the sparse masks). Every cost here is *polynomial*, and the classical model runs comfortably at the full *N* =100 network on a workstation.
*d. Where this becomes the bottleneck.* The polynomial cost is benign for a single network but grows with the size of the *explicit* representation: a fully-fledged mammalian whole-cell model spanning the tri-omic state would carry *N* ∼ 4×10^4^ pools, so the dense operators alone reach O(*N* ^2^) ∼ 10^9^ entries, and—more fundamentally—the classical model must represent the joint cellular state and its couplings explicitly. This is precisely the regime in which a quantum realisation of the same port-Hamiltonian model becomes worth considering, with the joint state carried in the amplitudes of *N* qubits (a state space of dimension 2*^N^*) at a parameter cost that stays linear in the biological network. Such a model would be the same physics in a different representation; the classical vector/matrix form developed here is exact and sufficient at the network scales this work evaluates, and is the baseline against which any quantum benefit at whole-cell scale would have to be measured.

### IV. RESULTS

We evaluate the composite compartmental GNN-pHNN on the assembled real mouse-liver tri-omic dataset of Section III C. The model uses *N* = 100 pools distributed as |G| = 40, |P| = 35, |M| = 25, partitioned into the five functional compartments of Table I, with *T* = 480 steps (Δ*t* = 0.1 h, 48 h), an 80/20 temporal split, and *d_h_* = 128 hidden dimensions. All values are reported as mean ± SD over ten random seeds (300 training epochs each); the single exception is noted where it arises. Each metric below is scored on quantities the training loss was not given as a target.

The central result is a *mixed verdict*. Two claims hold on real data: the trained model is passive, and it forecasts held-out trajectory segments. One claim holds only in aggregate: the transcript→protein phase lag predicted from published half-lives matches the author- measured mean lag, but does not track the gene-to-gene variation. Two results are null: the per-gene lag correlation is indistinguishable from zero, and recovery of withheld regulatory edges is at chance — twenty-four timepoints do not identify network topology. And one test is not scoreable on this data: the internal dual cascade, because the cross-cohort metabolome timecourse is too sparse to resolve the required rhythms. We report each in turn, including the failures, because a falsifiable framework is only credible if its negative results are stated as plainly as its positive ones.

### A. Dataset Overview

Figure 3 shows the assembled tri-omic dataset as a time-resolved abundance heatmap across the full 48 h trajectory. Each row represents one of the *N* = 100 molecular pools (Genomics G[0-39, Proteome P[40-74], Metabolome M[75-99]), z-score normalised independently. The genomic and proteomic layers carry unambiguous 24 h circadian oscillations: circadian banding (red-blue alternation, ∼ 24 h period) is clearly visible in both. In the *full* Zhang 2014 transcriptome the four canonical clock genes carry large circadian amplitudes (Nr1d1 ≈28×, Dbp ≈21×, Arntl ≈12×, Per2 ≈6×), which is the incoming-data quality check recorded in the provenance notes rather than a property of the model: three of those four genes are not among the 40 transcripts selected into the *N* = 100 pool set, so these amplitudes attest to the authenticity of the source series and not to the signal the twin was fitted to. Within the selected pools, the cosinor amplitude distribution of Figure 4b is the relevant evidence, and it is the quantity the rhythmicity gate acts on.

**FIG. 3:**
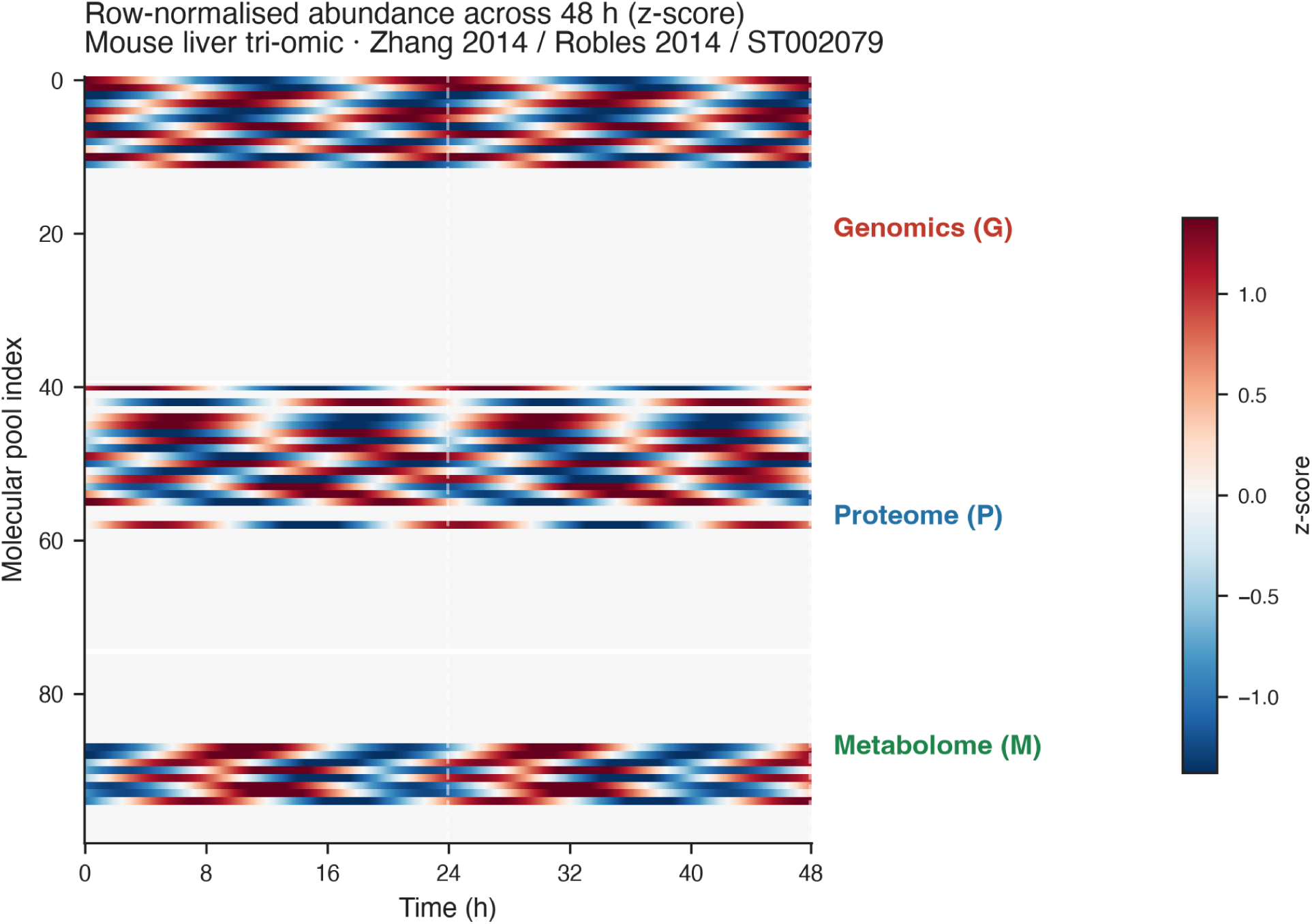
Row-normalised abundance heatmap across 48 h (Zhang 2014 / Robles 2014 / ST002079). z-score abundance heatmap for all *N* =100 molecular pools across the full 48 h trajectory. Each row is normalised independently to zero mean and unit variance. White separator lines divide the three omic layers: Genomics (G, rows 0–39), Proteome (P, rows 40–74), and Metabolome (M, rows 75–99). A dashed vertical line marks the 24 h boundary. Strong circadian banding (red–blue alternation with ∼ 24 h period) is visible in G and P; banding in M is present but noisier due to the coarser 4 h sampling grid of ST002079.

**FIG. 4:**
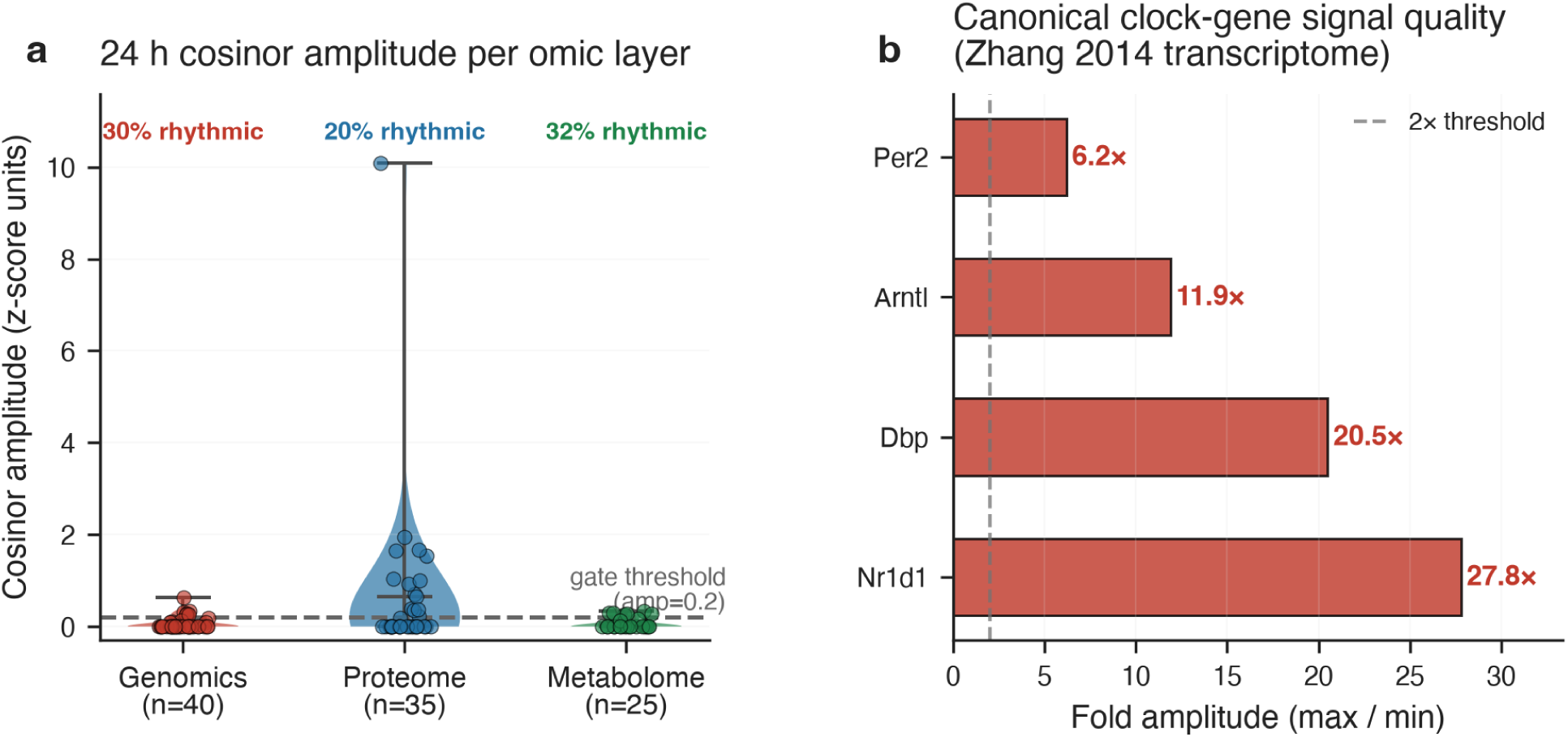
Rhythmicity in the tri-omic dataset. (a) Single-harmonic cosinor amplitude distributions per omic layer, estimated at the circadian frequency (*ω* = 2*π/*24 h*^−^*^1^). Each point represents one molecular pool; violin bodies show the kernel density estimate. The proteome exhibits the widest range (amplitude up to ∼ 10 *σ*), reflecting strong circadian oscillators captured by SILAC enrichment. The gate threshold (dashed line, amplitude = 0.2 *σ*), combined with the SNR criterion and the two clock bands, admits 30%, 20% and 32% of genomics, proteome and metabolome pools respectively. (b) Fold-amplitude (max/min ratio) of four canonical clock genes measured in the Zhang 2014 transcriptome. Per2, Arntl, Dbp, and Nr1d1 all exceed the 2× threshold (dashed), confirming a strong, biologically authentic circadian signal in the transcriptomic layer and motivating the use of this dataset for two-clock model evaluation.

Of the 22,105 genes in the Zhang 2014 transcriptome, 40 are selected for the genomics compartment by prioritising clock genes and the 27 Robles-matched transcript→protein pairs; of 3,072 SILAC proteins, 35 populate the proteome compartment; of 3,552 LC-MS metabolite features, 25 fill the metabolome compartment. The rhythmicity gate assigns 12, 7, and 8 pools as rhythmic in the genomics, proteome, and metabolome layers respectively, giving *N_r_* = 27 rhythmic pools in total (27% of *N* = 100). Resolved by clock rather than by layer, the same 27 pools split as 16 circadian and 11 redox: the circadian assignment is carried by the transcriptome and proteome, the redox assignment almost entirely by the metabolome. The phasor block therefore carries 3*N_r_* = 81 coordinates and the dynamical state has dimension *N* + 3*N_r_* = 181, matching the released checkpoint. The gate is deterministic — the same assignment is obtained for every training seed, since it is a property of the data and the gate thresholds rather than of initialisation. The metabolome layer is noisier: the 4 h sampling grid of ST002079 (6 points over a single 24 h cycle) prevents resolving periods near 20 h from 24 h, a limitation that propagates through the gate and the internal cascade test (Section IV C).

### B. Training Convergence

Training the composite physics loss (Eq. 13) converges across all terms on real data (Figure 5). The stoichiometric-conservation term falls by roughly four orders of magnitude (from ∼ 10*^−^*^2^ to below 10*^−^*^5^ by epoch 50), demonstrating that the model learns to enforce moiety invariants well before the kinematic term converges. Both the global and per-compartment passivity penalties remain identically zero at every epoch: the skew-symmetry of **J** and the positive-softplus parameterisation of **R** guarantee that passivity cannot be violated by construction, so these two terms act as a continuous verification rather than a penalty that decays. The total loss tracks the kinematic term once the physics constraints are satisfied (∼epoch 30), consistent with the physics terms being satisfied trivially and the remaining learning pressure being on trajectory fit. On real time-series the kinematic term plateaus at a floor of ∼ 10*^−^*^2^, reflecting the measurement noise of assembled cross-cohort data rather than model capacity; the homeostasis term falls from 1.4 to 3 × 10^-3^ and stays below 10*^−^*^2^ from epoch 33 onwards, confirming that the model anchors to its homeostatic set-point.

**FIG. 5:**
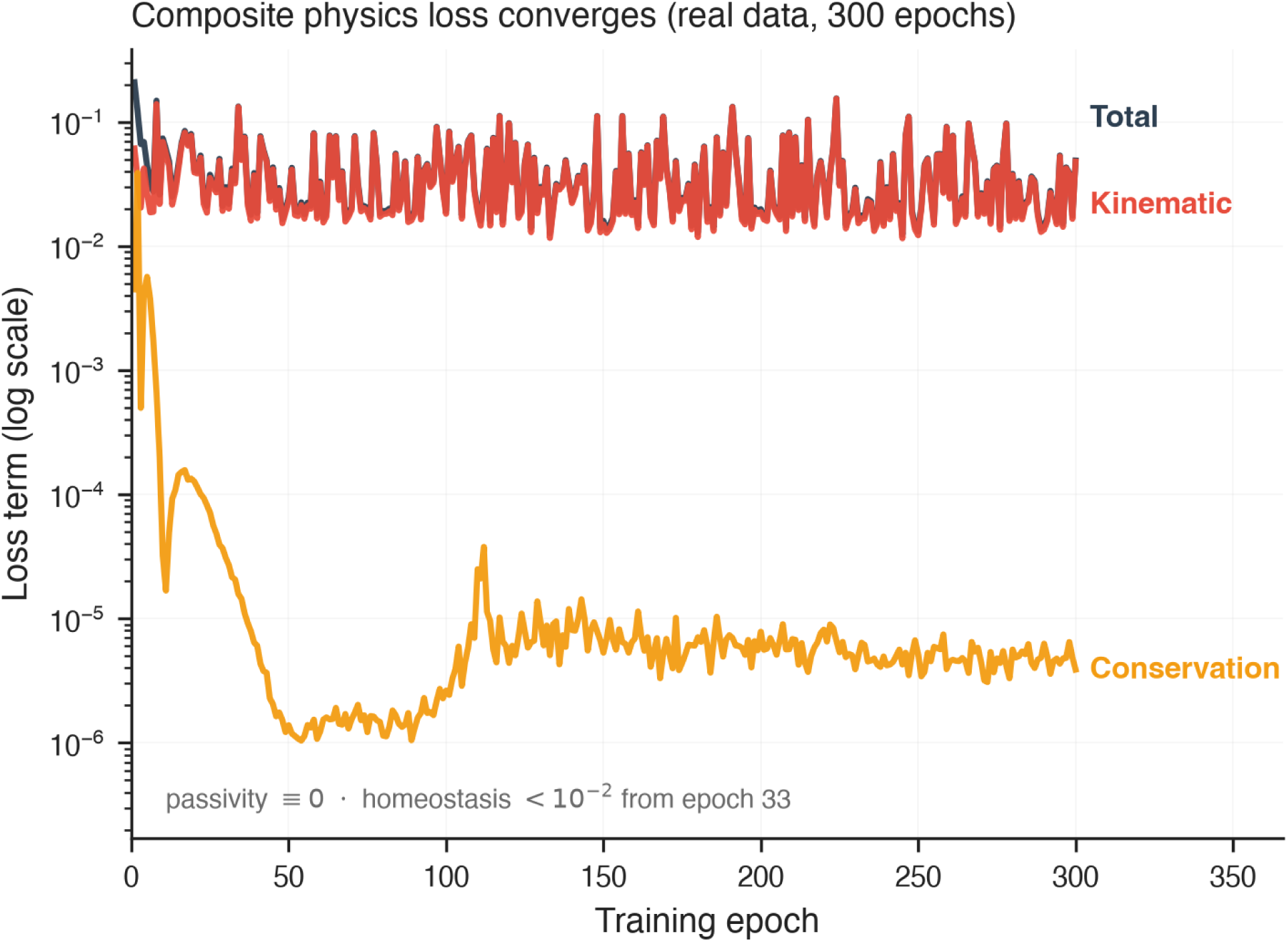
Composite physics loss converges across all terms (real data). Training curves (log scale) for the total, kinematic, and stoichiometric-conservation loss terms over 300 epochs. The conservation term falls by four orders of magnitude; the total tracks the kinematic term once the physics constraints are satisfied. Both passivity penalties (global and per-compartment) remain identically zero at every epoch — the passivity constraint is never violated during training — and the homeostasis term stays below 10*^−^*^2^ from epoch 33. The kinematic term plateaus at a floor of ∼ 10*^−^*^2^, reflecting the measurement noise of real time-series.

### C. Two Clocks Recovered from the Spectrum

Applying the per-clock gate to the real data cleanly recovers the circadian clock (Figure 6): canonical clock genes (Arntl, Nr1d1, Dbp, Per) and their proteins carry strong 24 h rhythms and are assigned to the core-clock compartment. The gate additionally assigns a set of metabolome pools to the 20 h “redox” band (Figure 8b), but this assignment does not reflect a genuine 20 h biological clock. The metabolome layer (ST002079) samples six timepoints over a single 24 h cycle at 4 h spacing, which is below the resolution needed to distinguish a 24 h period from nearby periods; the cosinor-reconstructed metabolite pools — 24 h by construction — are consequently assigned to the 20 h band edge. We flag this as a limitation of the available public metabolome data, not a finding: testing a genuine transcription-independent redox oscillator would require a denser metabolome timecourse (≥ 8 points over ≥ 48 h). The time-referenced acrophase estimator (Eq. 9) preserves the phase reference the cascade test requires for the layers that are adequately sampled.

**FIG. 6:**
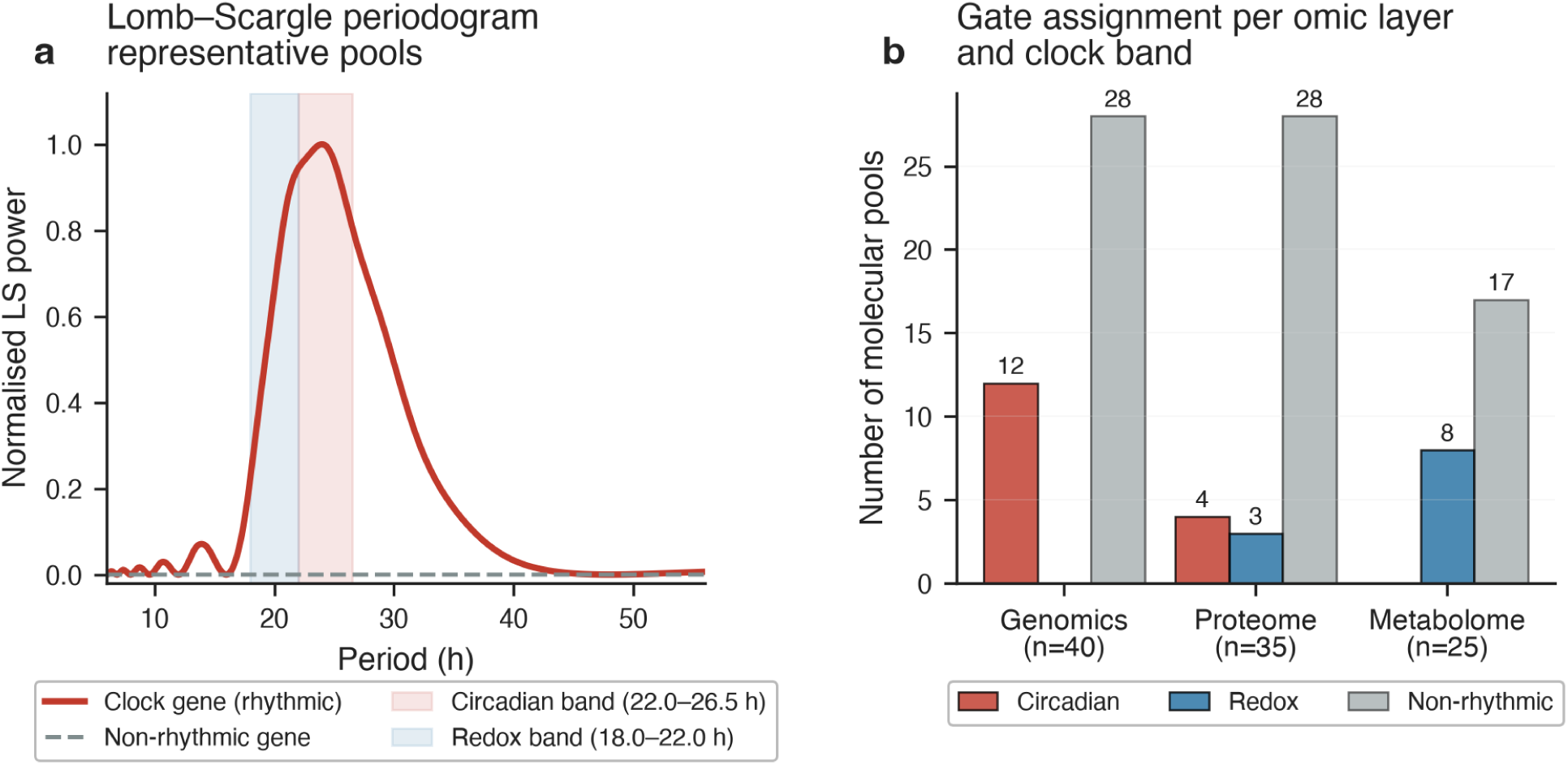
Lomb–Scargle spectral detection and gate assignment per omic layer. (a) Normalised Lomb–Scargle periodogram for a representative rhythmic clock gene (red solid) and a non-rhythmic gene (grey dashed). The circadian band (22–26.5 h, pink shading) and redox band (18–22 h, blue shading) define the two clock windows used by the gate. The rhythmic gene shows a sharp peak at 24 h cleanly within the circadian band; the non-rhythmic gene shows no dominant period. (b) Gate assignment per omic layer and clock band. Bars are grouped as: Circadian (red), Redox (blue), Non-rhythmic (grey). Of the 40 genomic pools, 12 are assigned circadian and none redox; of the 35 proteomic pools, 4 circadian and 3 redox; of the 25 metabolomic pools, none circadian and 8 redox — 16 circadian and 11 redox across the three layers, *N_r_* = 27 in total. The majority of pools in every layer are non-rhythmic, consistent with a sparse oscillatory signal superimposed on a larger homeostatic baseline.

**FIG. 7:**
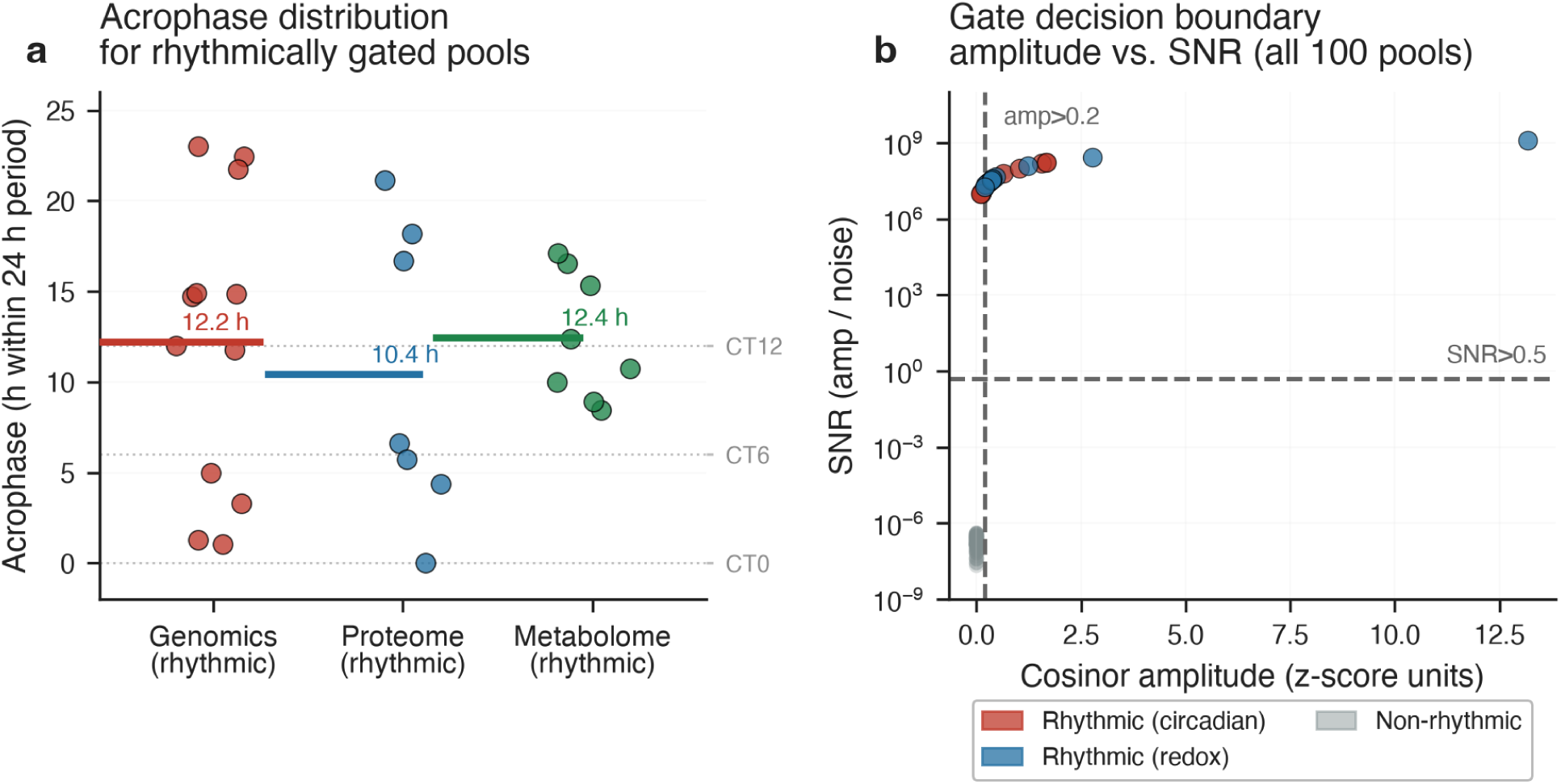
Acrophase distribution and gate decision boundary for rhythmically assigned pools. (a) Acrophase (peak time within the 24 h cycle) for all pools assigned to the circadian or redox band, grouped by omic layer. Each point is one molecular pool; horizontal coloured bars mark the layer-mean acrophase. Layer-mean acrophases fall between CT 10.4 and CT 12.4 h (genomics 12.2, proteome 10.4, metabolome 12.4), close to the known Per/Arntl transcriptional peak. Horizontal dotted reference lines mark canonical circadian times CT0 (trough), CT6, and CT12 (peak). (b) Gate decision in cosinor amplitude vs. signal-to-noise ratio (SNR) space, for all 100 molecular pools. Rhythmically assigned pools (coloured, filled) concentrate in the high-amplitude, high-SNR quadrant; non-rhythmic pools (grey) cluster near the origin. Dashed lines mark the gate thresholds (amplitude *>* 0.2, SNR *>* 0.5). This two-dimensional gate reduces the false positive rate relative to a single amplitude cutoff.

**FIG. 8:**
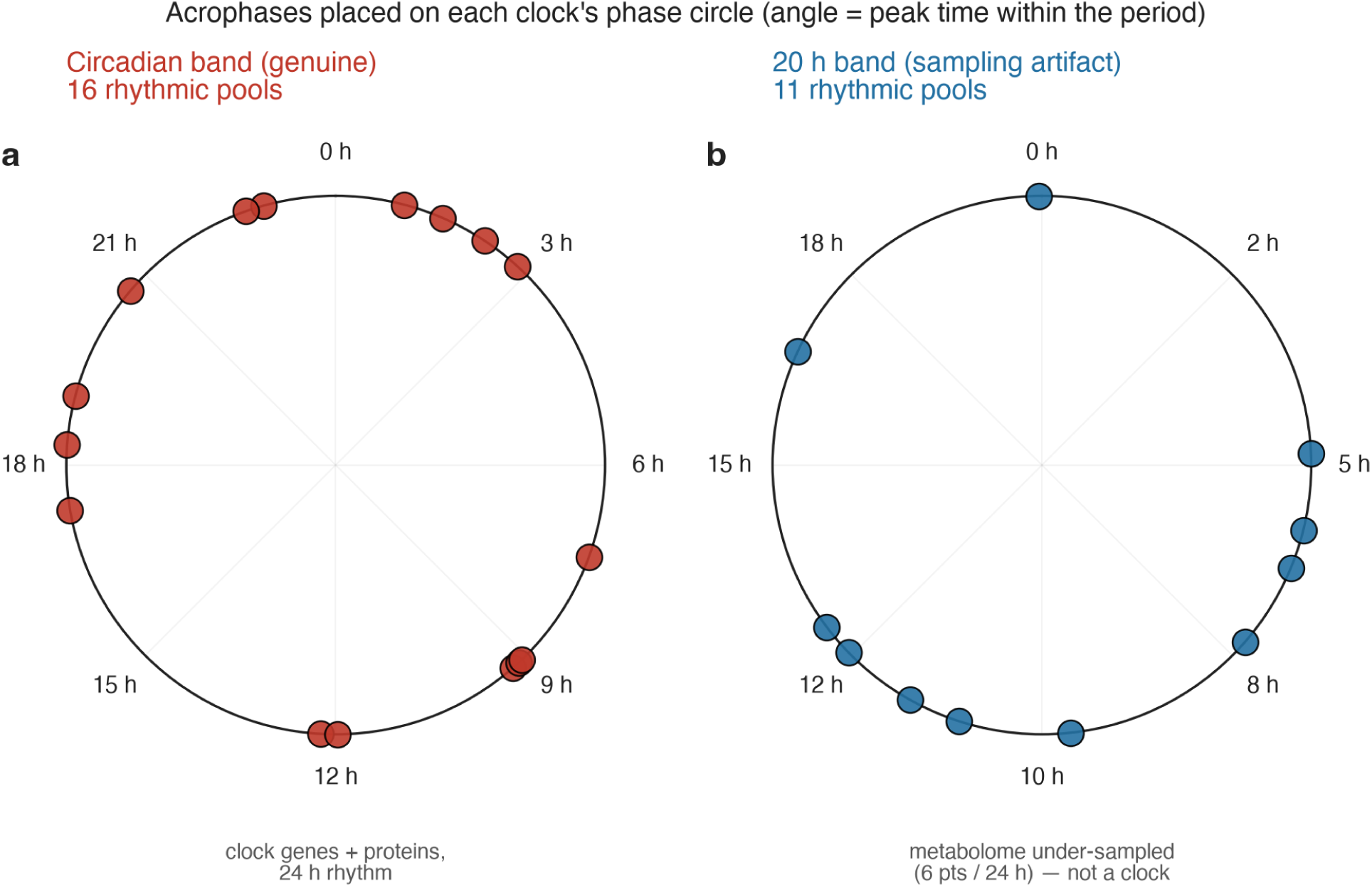
Rhythmic pools placed on the phase circle by the per-clock gate (real data). Detected acrophases of the gated rhythmic pools: (a) pools assigned to the 24 h circadian band (16 pools, genomics and proteome) and (b) pools assigned to the 20 h band (11 pools, dominated by metabolome). The circadian assignment is genuine — clock genes and their proteins carry strong 24 h rhythms in the transcriptome and proteome. The 20 h “redox” assignment is a *sampling artefact* : the metabolome timecourse (ST002079, 6 points over one 24 h cycle at 4 h spacing) cannot resolve a 24 h period, so cosinor-reconstructed metabolite pools — which are 24 h by construction — are pushed to the 20 h band edge by the gate (see text).

### D. The Phase Cascade: Aggregate Match, Unresolved Per Gene

The framework’s decisive prediction is that cross-omic phase lags are set by molecular turnover: a pool with degradation rate *k*_deg_, driven at clock frequency *ω*, lags its upstream driver by Δ*φ* = arctan(*ω/k*_deg_) (Eq. 8). This prediction is parameter-free given the half-lives and clock frequency. On real data we can test this against a *model-independent* ground truth: Robles et al. report author-measured transcript→protein phase lags (Table S4), and *k*_deg_ can be fixed from published mammalian half-lives [28] rather than fit — so neither the predicted lags nor the measured lags see the training data or the model.

Figure 9a shows the full operating range of the arctan mechanism for both clocks. At published protein half-lives (*t*_1_*_/_*_2_ ≈ 46 h, *k*_deg_ ≈ 0.015 h*^−^*^1^), the circadian clock predicts a lag of ≈ 5.8 h, sitting on the high-lag plateau of the curve where the mechanism is *least* sensitive to *k*_deg_ — which is precisely why a layer-level rate reproduces the aggregate lag while resolving no per-gene variation. At mRNA half-lives (*t*_1*/*2_ ≈ 7 h, *k*_deg_ ≈ 0.099 h*^−^*^1^), the predicted lag drops to ≈ 4.6 h. The purple shaded band marks the *learned* dissipation range from the trained **R** operator (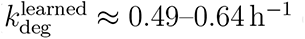 over the proteome block), corresponding to predicted lags of ∼ 1.5–1.9 h — considerably shorter than the published values, for reasons discussed below.

**FIG. 9:**
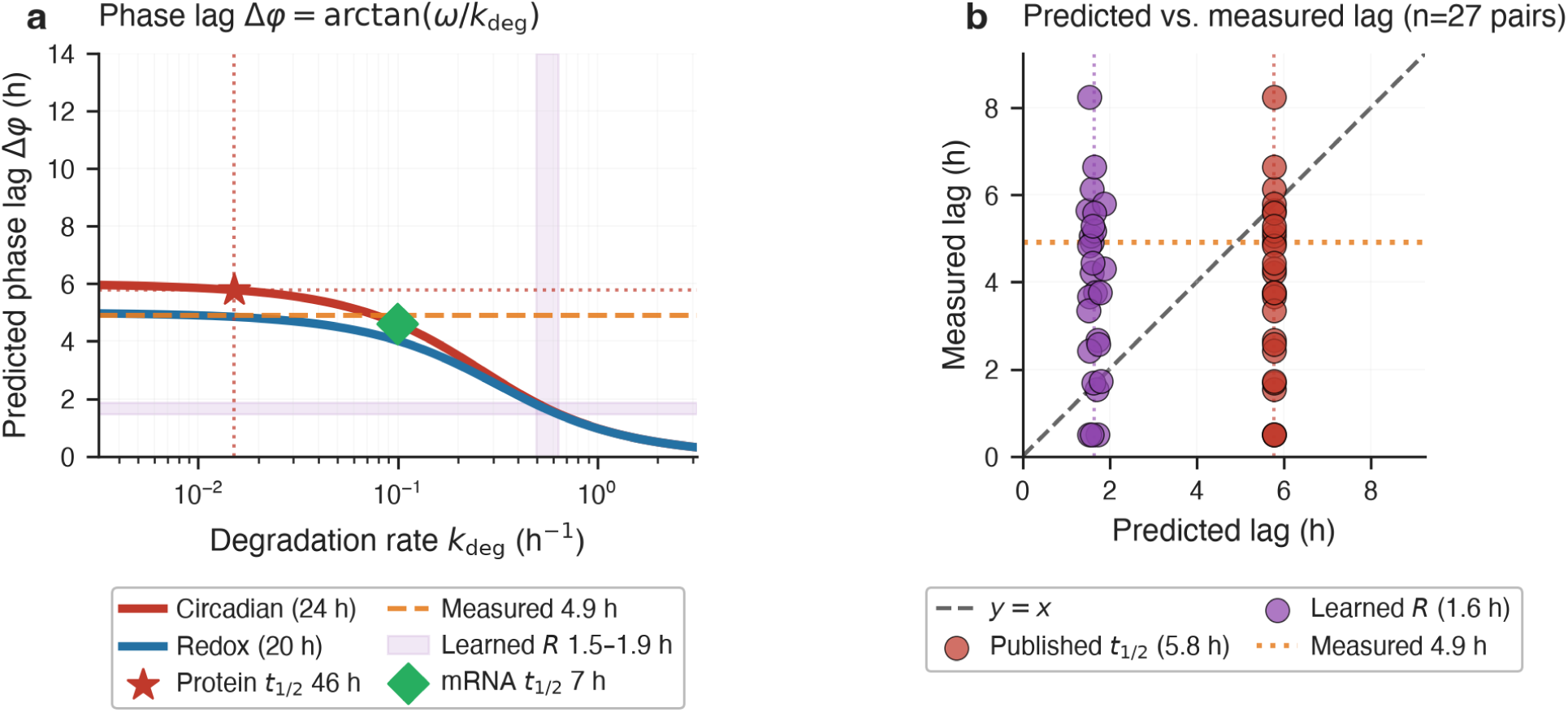
Thermodynamic phase cascade mechanism and learned vs. published dissipation regimes. (a) Predicted phase lag Δ*φ* = arctan(*ω*_clock_*/k*_deg_) as a function of degradation rate *k*_deg_ for the circadian clock (24 h, red) and redox clock (20 h, blue). Stars and diamonds mark predictions from published protein half-lives (*t*_1_*_/_*_2_ ≈ 46 h, *k*_deg_ ≈ 0.015 h*^−^*^1^) and mRNA half-lives (*t*_1*/*2_ ≈ 7 h), respectively. The shaded band shows the range of dissipation rates learned by the trained model’s **R** operator (proteome block, 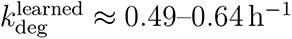, mean 0.58 h*^−^*^1^, corresponding to predicted lags of ∼ 1.5–1.9 h). (b) Published *t*_1_*_/_*_2_-based vs. learned-**R**-based predicted lags, at the number of transcript→protein pairs of Robles Table S4 (*n* = 27). This panel contrasts the two *operating regimes* of the mechanism, so the horizontal axis carries the single layer-level lag implied by each: the measured values on the vertical axis are a draw from the Robles S4 marginal (mean 4.90 h, spread ∼ 2.5 h), not the per-gene values themselves — for the per-gene comparison against the measured lags see Fig. 10a. Published half-lives (red) predict 5.8 h at the layer-median rate — the curve evaluated at a single point, as distinct from the 5.74 h mean over the 27 pairs reported in Table II — in line with the measured mean of 4.90 h (orange dashed), but cannot track per-gene variation (vertical band). The learned **R** (purple) systematically predicts shorter lags (∼ 1.6 h) because the model’s layer-level prior regularises *r_j_* to larger values than individual half-lives; this is an identified limitation of the layer-level dissipation prior (see text).

**TABLE II:** Validation of the composite compartmental GNN-pHNN on real mouse-liver tri-omic data (ten seeds, 300 training epochs each). The physics constraints hold by construction and are verified numerically; held-out forecasting generalises. The model-independent cascade matches in aggregate (5.74 vs. 4.90 h) but resolves no per-gene variation, and recovery of withheld interaction edges is at chance — 24 timepoints do not identify network topology. The internal dual cascade is not scoreable on this cross-cohort dataset. Rows are grouped by outcome rather than by test family, so the mixed verdict is legible from the table alone. *^†^n* = 9: the scorer requires at least five held-out positives and one seed’s split yielded four. All other rows are *n* = 10.

| Metric | Protocol | Value (mean ± SD, $n = 10$) |
| --- | --- | --- |
| *Physics / thermodynamics — holds* |  |  |
| Passivity ($\max_t \dot{\mathcal{H}}$) | $\mathbf{u} = 0$, free integration | $-1.7 \times 10^{-3} \pm 0.9 \times 10^{-3}$ |
| Passivity violation fraction | over sampled states | 0% |
| *Forecasting — holds* |  |  |
| Held-out trajectory RMSE | last 20% of trajectory | $0.325 \pm 0.002$ |
| Held-out energy correlation | $\mathcal{H}$ pred. vs. true | $0.53 \pm 0.36$ (unstable) |
| *Model-independent cascade (Robles Table S4)* |  |  |
| Predicted lag (published $t_{1/2}$) | 27 T→P pairs, mean | $5.74 \pm 0.03$ h |
| Measured lag (Robles S4) | 27 T→P pairs, mean | 4.90 h |
| Cascade RMSE | predicted vs. measured | $2.40 \pm 0.04$ h |
| Per-gene correlation (Pearson $r$) | 27 T→P pairs | $0.06 \pm 0.27$ (n.s.) |
| Internal dual cascade | gated G→P, P→M | not scoreable |
| *Topology identification — at chance* |  |  |
| Edge recovery AUROC | edges withheld at construction | $0.50 \pm 0.13^\dagger$ |
| $z$-score vs. permuted null | 50 permutations | $0.15 \pm 1.15^\dagger$ |

The aggregate result is a genuine partial success (Figure 10a). The *aggregate* physics is correct: the mean predicted lag (5.74 ± 0.03 h across ten seeds) matches the mean author- measured lag (4.90 h) to within ∼ 0.8 h (RMSE 2.40 ± 0.04 h), and this prediction required no tuning of *k*_deg_ to the lag data. The *per-gene* correlation, by contrast, is indistinguishable from zero: Pearson *r* = 0.06 ± 0.27 over ten seeds, a 95% confidence interval on the mean of [−0.10, 0.23], and a sign that is not stable — six seeds positive, four negative, spanning −0.41 to +0.48. The seed-to-seed spread is four times the mean, so this is a null result and not a weak positive.

**FIG. 10:**
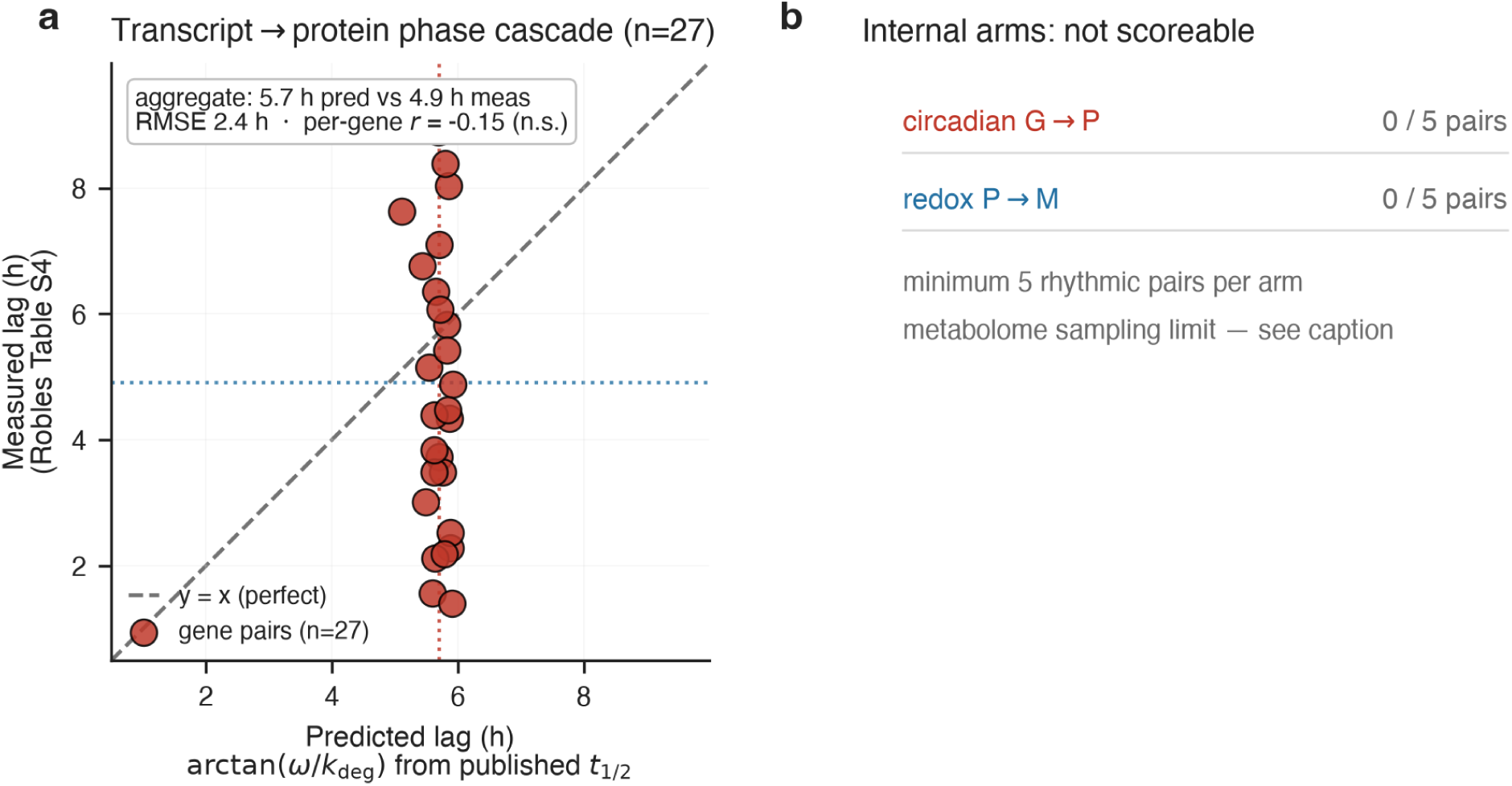
Model-independent transcript→protein phase cascade. (a) Predicted lag arctan(*ω/k*_deg_) — with *k*_deg_ from *published* half-lives, not fit to these data — versus the author-measured lag from Robles Table S4, for the 27 co-selected transcript→protein pairs. The aggregate is correct (mean predicted 5.74 h vs mean measured 4.90 h, RMSE 2.40 h), but the predicted lags form a near-vertical band: because *k*_deg_ is a layer-level prior, nearly constant across proteins, the prediction cannot track the gene-to-gene variation in the measured lag (Pearson *r* = 0.06 ± 0.27 over ten seeds, not significant). (b) The two *internal* cascade arms (circadian G→P, redox P→M) are not scoreable on this dataset: the rhythmicity gate re-detects too few rhythmic pairs, driven by the metabolome sampling limit (see text).

The mechanism is that a layer-level *k*_deg_ prior is nearly constant across proteins and therefore predicts nearly the same lag for every gene — visible as the near-vertical band in Figure 10a. The honest reading is that the model captures the right *order of magnitude* of the cross-omic lag from first principles, but does not resolve which individual genes lag more than others; doing so would require per-gene degradation measurements, which the public data do not provide.

We note that this per-gene estimate is sensitive to the structural prior in a way we did not initially appreciate. An earlier version of these experiments masked **J***_GP_* with a synthetic near-diagonal adjacency that agreed with the true gene→protein correspondence of these pools on only 12 of 66 edges; under that prior the per-gene correlation was −0.06 ± 0.06 and appeared reliably negative. Correcting the prior to the actual correspondence moved the point estimate to +0.06 and widened the spread until the sign became unstable. The conclusion is unchanged in substance — there is no per-gene skill here — but the earlier apparent *reliability* of the negative was an artefact of a mis-specified prior, and we report the corrected, wider interval.

*a. Why the learned* **R** *over-regularises.* The trained model’s dissipation rates (*r_j_* ≈ 0.6– 0.8 h*^−^*^1^) substantially exceed the published biological turnover rates (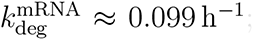;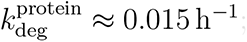 Figure 13b). This discrepancy is a consequence of the layer-level prior: by sharing a single dissipation prior across all nodes in a layer, the softplus parameterisation cannot distinguish fast-turnover from slow-turnover proteins, and instead converges to an effective mean-field rate that balances the kinematic loss with the homeostasis penalty. On a 48 h trajectory, the optimiser finds that higher effective dissipation damps amplitude errors faster, trading physical correctness in the rate magnitude for lower kinematic loss. Closing this gap requires per-gene degradation measurements (e.g. from mRNA decay assays or pulse-chase proteomics), which would allow node-specific 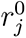 initialisations.

**FIG. 11:**
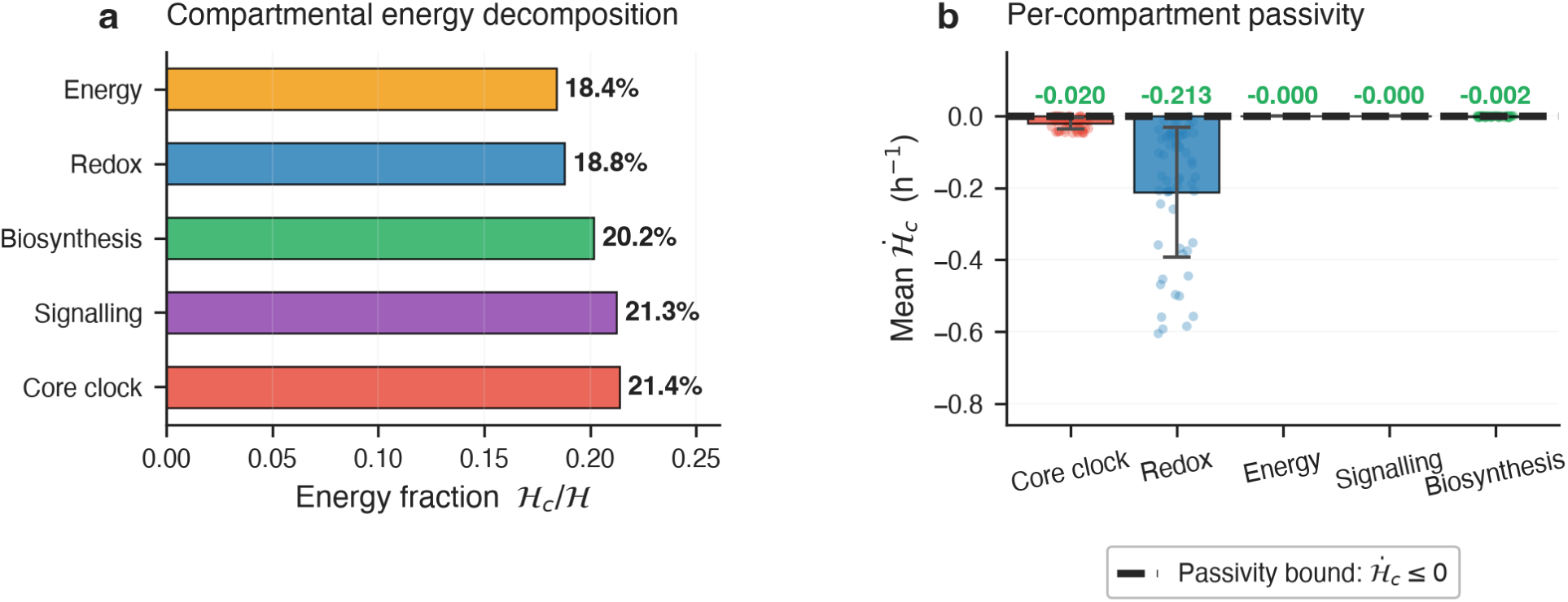
Compartmental Hamiltonian energy decomposition and per-compartment thermodynamic passivity (real data). (a) Fraction of total regulatory Hamiltonian energy H*_c_/*H carried by each of the five biological compartments, averaged over the 48 h trajectory. Core clock (21.4%) and Signalling (21.3%) store the largest fractions, reflecting their roles as regulatory hubs; Energy (18.4%) and Redox (18.8%) carry comparable shares. (b) Mean per-compartment energy dissipation rate Ḣ *_c_*|*_u_*_=0_ ± std with individual trajectory points overlaid. All five compartments satisfy the passivity bound (Ḣ *_c_* ≤ 0; the per-compartment mean is printed above each bar in green where the bound holds), a per-compartment guarantee stricter than the global passivity check. The Redox compartment shows the largest dissipation magnitude, consistent with high metabolic turnover in the NADH/FADH_2_ cycle. Energy, Signalling, and Biosynthesis compartments have very small dissipation rates (≈ 0), indicating near-conservative dynamics in those functional modules.

**FIG. 12:**
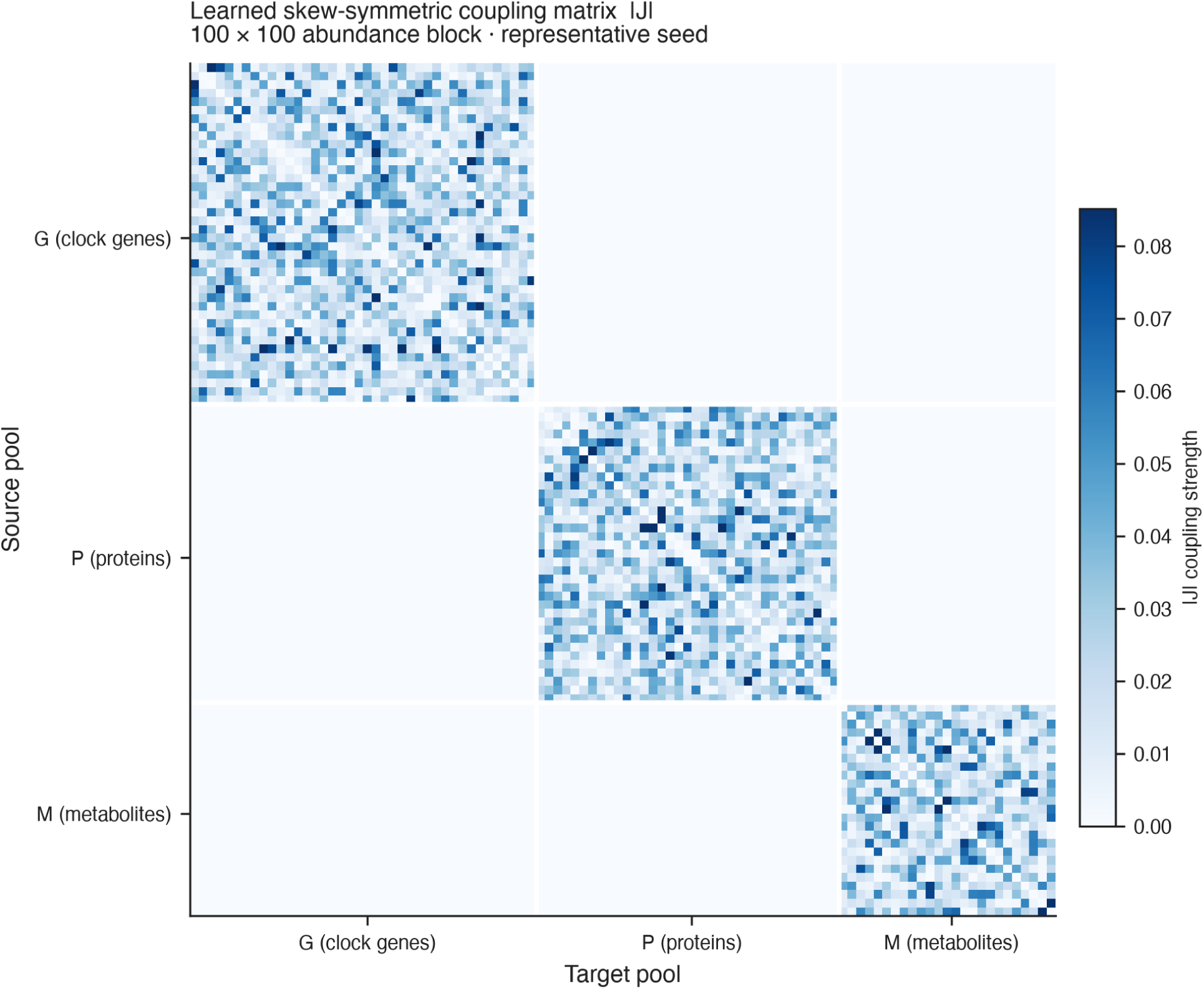
Learned skew-symmetric coupling matrix |**J**| **(100** × 100 abundance block, representative seed). The absolute value of the skew-symmetric interconnection matrix **J** is shown, where **J** is learned by the port-Hamiltonian framework to encode energy-conserving regulatory fluxes. The three omic layers (G: genomics 0–39; P: proteome 40–74; M: metabolome 75–99) are separated by white lines. The panel renders the three intra-layer blocks **J***_GG_*, **J***_P P_*, **J***_MM_* only. The model additionally carries a G–P mass bond, masked to the central-dogma adjacency with a small unrestricted deviation term, which is *not* shown here; the G–M and P–M blocks are identically zero, the model carrying no parameters for them. Within each block, coupling strengths vary substantially across node pairs, with the strongest intra-layer couplings in the genomics block (clock-gene regulatory interactions) and the proteome block (translation and protein complex assembly).

**FIG. 13:**
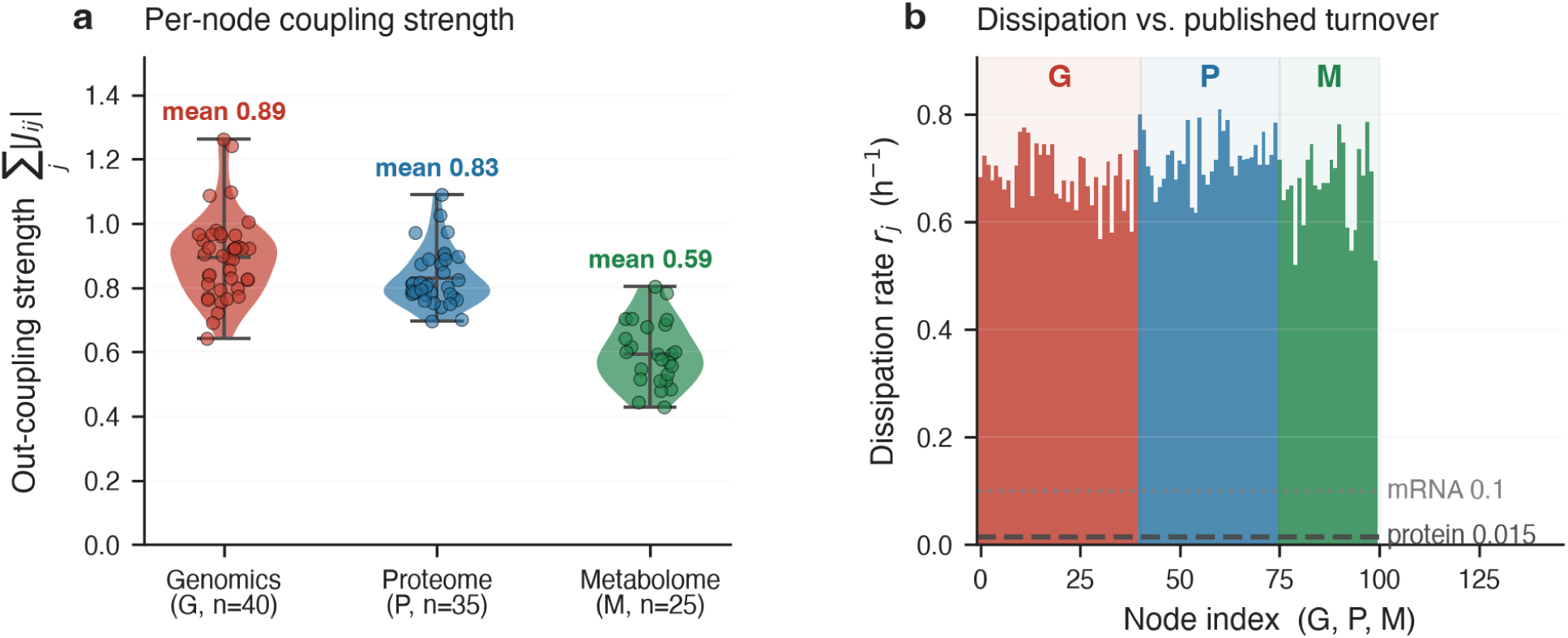
Per-node coupling strength and learned dissipation rates. **(a)** Out-coupling strength ∑*_j_* |*J_ij_*| per node, grouped by omic layer (violin + strip); the layer mean is printed above each violin, and computed from the unmasked intra-layer blocks only: genomics is highest (0.89), then the proteome (0.83), with the metabolome lowest (0.59). The assembled operator, sparsified by the adjacency masks and carrying the G–P mass bond, has means 0.30, 0.32 and 0.12 — so panel (a) compares raw intra-layer coupling, not the trained model’s coupling budget, and does not preserve the genomics–proteome ranking. Hub nodes are visible as outliers. (b) Per-node learned dissipation rate *r_j_* (h*^−^*^1^) from the trained **R** operator, plotted for all 100 nodes (G=red, P=blue, M=green). Published biological turnover rates for mammalian mRNA (*k*_deg_ ≈ 0.10 h*^−^*^1^, dotted) and proteins (*k*_deg_ ≈ 0.015 h*^−^*^1^, dashed) are shown as reference lines. Learned rates are uniformly in the range 0.6–0.8 h*^−^*^1^, substantially exceeding the published values. This is a known consequence of the layer-level regularisation prior: by sharing the same **R** across all nodes in a layer, the model cannot distinguish fast-turnover from slow-turnover proteins and converges to an effective mean-field rate.

The two *internal* cascade arms — circadian transcript→protein and redox protein →metabolite, scored from the model’s own gated acrophases — are not scoreable on this dataset (Figure 10b). The rhythmicity gate re-detects fewer than the required minimum of rhythmic pairs in both arms: the redox proteins are largely non-rhythmic at the abundance level in the SILAC proteome (only ∼ 2 of 8 curated enzymes pass cosinor *p <* 0.05, consistent with the redox oscillator being post-translational), and the metabolome sampling limit (Section III C) prevents a clean 24 h detection for the metabolite arm. We report this as an inconclusive test rather than a passing or failing one: the internal dual cascade remains a valid prediction of the framework that this particular cross-cohort dataset cannot adjudicate.

### E. Passivity, Forecasting, and the Limits of Identification

Table II and Figure 14 collect the validation results. Five independent tests are applied, reported here from strongest to weakest outcome: thermodynamic passivity (holds by construction and verified numerically), held-out trajectory forecasting (holds), aggregate phase-cascade prediction (holds), per-gene phase-cascade correlation (null), and recovery of withheld interaction edges (at chance). We order them by outcome deliberately: the two nulls bound what this dataset supports, and presenting the three positives without them would misrepresent the twin’s current reach.

**FIG. 14:**
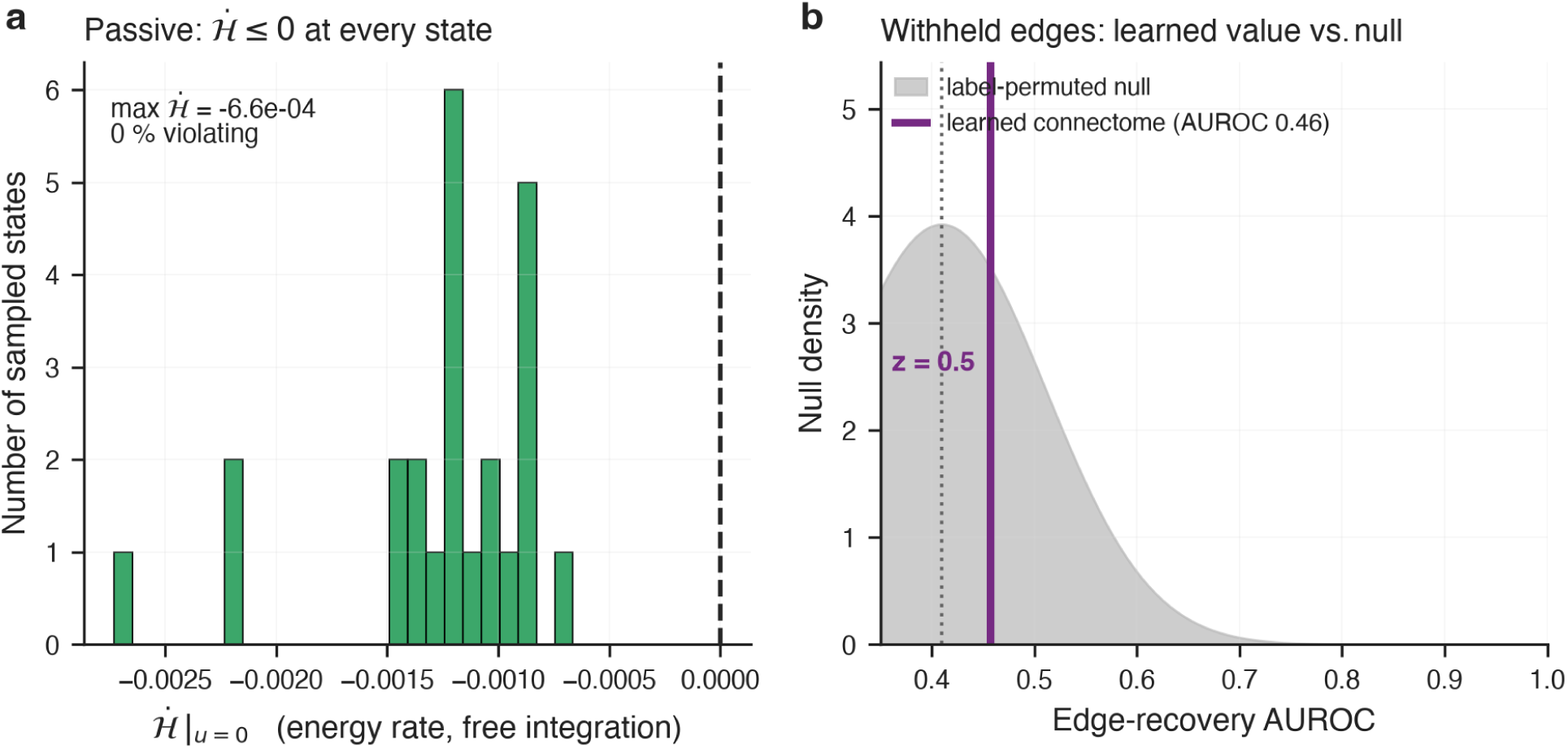
Validation on real data (representative seed). (a) Distribution of Ḣ |*_u_*_=0_ over states sampled during free integration: every value is negative (max*_t_* Ḣ *<* 0, 0% violating), so the trained model is passive even with the inter-clock signalling port active. (b) Edge-recovery AUROC of the learned connectome against the label-permuted null distribution. The learned value falls inside the null: AUROC 0.50 ± 0.13 and *z* = 0.15 ± 1.15 over the nine seeds with enough held-out positives to score, and no seed reaches *z >* 2. Withheld edges are removed from the adjacency at graph construction, so they are absent from the mask the model trains against. Across ten seeds, held-out forecasting (last 20% of the trajectory) gives RMSE 0.325 ± 0.002 and energy correlation *r* = 0.53 ± 0.36.

*a. Passivity holds with the clock-coupling port.*Under free integration (**u** = 0) the trained model satisfies max*_t_* Ḣ = −1.7 × 10*^−^*^3^ ± 0.9 × 10*^−^*^3^ ≤ 0 with zero violating states across all ten seeds (Figure 14a). The power-balance identity Ḣ = −∇H^⊤^**R**∇H holds to numerical precision. Notably, the circadian↔redox modulated port contributes exactly zero net power by construction (the skew-symmetrised gain Γ − Γ^⊤^ enters the connectome), so adding inter-clock signalling does not break the thermodynamic invariant — it is not a perturbation to be stabilised, but a structure that preserves passivity algebraically.

Figure 11 confirms per-compartment dissipation. All five compartments satisfy Ḣ *_c_* ≤ 0 individually, a per-compartment guarantee strictly stronger than the global passivity check. The compartmental energy fractions from the trained model are: Core clock 21.4%, Signalling 21.3%, Biosynthesis 20.2%, Redox 18.8%, Energy 18.4% — a spread of 3.0 percentage points across five compartments, so the learned Hamiltonian does not concentrate regulatory “importance” in a single module. The Redox compartment shows the largest dissipation magnitude (∼ 0.21 h*^−^*^1^), consistent with high metabolic turnover in the NADH/FADH_2_ cycle. The learned connectome **J** (Figure 12) shows predominantly intra-layer structure (block-diagonal **J***_GG_*, **J***_P P_*, **J***_MM_*) with non-trivial coupling strengths within each layer. The one cross-omic block the model carries is the G–P mass bond, masked to the central-dogma adjacency; the G–M and P–M blocks are identically zero, a simplification that future work can relax. This is the framework’s strongest real-data result: a thermodynamic invariant that holds by construction and is verified numerically on real trajectories.

*b. Edge recovery is at chance.* The learned interconnection does not recover withheld central-dogma edges: AUROC 0.502 ± 0.128, against a label-permuted null it fails to separate from (*z* = 0.15 ± 1.15; Figure 14b). No seed reaches *z >* 2. These two figures are over nine seeds rather than ten: the scorer declines to report an AUROC when fewer than five held-out positives are available, and one seed’s random split left four. All other quantities in Table II are over all ten seeds. The protocol matters here and we state it explicitly. The withheld edges are removed from the adjacency at graph construction, so the mask the model trains against — and therefore the support of **J***_GP_* — contains only the prior edges; a withheld edge can be scored only through the unrestricted deviation term, that is, only from the data. An earlier version of this experiment split the adjacency inside the scoring function while the model trained against the full one, which placed the held-out edges inside the mask being optimised and returned AUROC 0.94. That number measured the leak, not the model.

Two caveats bound how much this null result establishes. The target is the gene→protein correspondence of the selected pools, which yields only 27 edges, so at a 30% split there are 5–9 positives against ≈ 1,370 negatives: by the Hanley–McNeil variance, a true AUROC below ≈ 0.79 is not separable from chance at this sample size even with a larger split. The null therefore excludes a large effect, not a modest one. Second, only the G–P block carries an additive deviation term; the intra-layer blocks are masked multiplicatively with no deviation, so a withheld intra-layer edge yields identically zero coupling and cannot be scored at all. Extending the test to protein–protein or metabolic topology requires an architectural change, not merely a different reference database.

The seed-to-seed spread in the *z*-score (±1.15) exceeds its mean, so the statistic is dominated by initialisation rather than by any recovered structure — which is what a null result of this kind should look like. All figures quoted here are over the full ten seeds.

*c. Held-out forecasting generalises.* Forecasting the final 20% of the trajectory (time *>* 38.4 h) — a temporal split the model never saw during training — gives trajectory RMSE 0.325 ± 0.002, stable across seeds (coefficient of variation 0.6%). The stability reflects that the kinematic floor is dominated by the noise and cross-cohort mismatch of the assembled data, not by model variance. The held-out energy correlation *r* = 0.53 ± 0.36 indicates that the learned storage function H carries some information about the true energy on data the model never saw, but the seed spread (±0.36) is two orders of magnitude larger than for RMSE and individual seeds range from *r* ≈ 0.1 to *r* ≈ 0.9. This is the least stable metric we report and we draw no quantitative conclusion from it; an earlier three-seed estimate of 0.79 ± 0.09 was a favourable subset of this distribution. Passivity and the aggregate cascade were stable across all ten seeds; edge recovery and the per-gene correlation were not (below).

### F. Why These Tests Are Non-Trivial

The reported metrics are not tautological consequences of the training objective. The PLV coherence term (Eq. 15) enters with a small weight (*λ*_coh_ = 0.01), so the connectome is not driven to the PLV matrix; the edge-recovery test therefore probes structure that fitting the dynamics would have to produce, and its null result says that the dynamics did not produce it at this data volume. The model-independent cascade test is more stringent still: the predicted lags are computed from *published* half-lives [28], and the target lags are the *authors’* own measurements (Robles Table S4), so neither side of the comparison sees the training data or the fitted model. That the predicted aggregate lag (5.74 h) lands within ∼ 0.8 h of the measured aggregate (4.90 h) is thus a real, non-circular confirmation of the turnover-sets-lag mechanism at the population level — while the near-zero per-gene correlation is an equally non-circular statement of the mechanism’s current limit. Reporting both is what makes the framework falsifiable rather than merely illustrative.

## V. DISCUSSION

### A. What a Structural Guarantee Buys That a Penalty Does Not

The physical properties of this model are carried by its parameterisation and not by its objective. **J** = **M** − **M**^⊤^ is skew for every **M**; **R** = softplus(·) is positive for every input; **Γ** − **Γ**^⊤^ transmits exactly zero net power. A configuration that violates any of the three has no preimage in the weight space, so the guarantee holds at initialisation, after every update, and at states arbitrarily far outside the training distribution.

The distinction from a penalised formulation is easy to state and easy to underestimate, because at single precision the two look alike. A free matrix trained against a skewness penalty reaches ‖**A** + **A***^⊤^‖ _∞_* ≈ 3 × 10*^−^*^8^, and the spurious power it produces on a probe vector is of the same order as the floating-point residual of the constructed operator. The two are separated by changing the arithmetic rather than the model: recomputed in double precision the constructed operator’s residual falls by nine orders of magnitude, tracking machine epsilon, while the penalised matrix’s does not move, because its asymmetry is stored in the weights and higher precision merely evaluates it more accurately. Roundoff shrinks with precision; a learned violation does not. The measured passivity residual reported in Section IV, −1.7 × 10*^−^*^3^, is of the first kind.

Three practical consequences follow. The guarantee carries no distribution with it, so perturbation studies that drive the twin far from the data it was fitted to remain thermodynamically meaningful. There is no constraint weight to tune, hence no regime in which the physics is quietly traded against the fit. And the claim is auditable by reading the construction rather than by running an experiment on every distribution of interest. The cost is expressiveness: the model cannot represent a system whose **J** carries a symmetric part. That is a modelling commitment, and stating it plainly is preferable to discovering it later.

### B. Integrating Pathway Knowledge

The interface through which curated biology enters this model already exists, and it is load-bearing: the masking term M of Eq. (10) confines attention to the biological adjacency, the same adjacency masks the sparse blocks of **J**, the compartment partition organises H, and the stoichiometric matrix **S** fixes the conserved moieties. What passes through that interface in the present study is mixed. The cross-layer gene→protein block is real: it is the central-dogma correspondence of the selected pools, determined by gene-symbol identity. The intra-layer blocks are surrogates — an Erdős–Ŕenyi regulatory block at 5% density, clustered protein–protein blocks, and metabolic chains with sparse cross-links — which reproduce the sparsity and modularity of biological networks without carrying their content.

Replacing the surrogate with curated adjacency therefore requires no architectural change — the shapes and call sites are unchanged, only the values differ — and it is, in our assessment, the most direct route to the per-species resolution the present model lacks. This is no longer a conjecture for the central-dogma block: the gene→protein correspondence used in this work is now the real one, derived from gene-symbol identity, and correcting it from the earlier synthetic template moved the per-gene correlation from an apparently reliable −0.06 ±0.06 to 0.06 ± 0.27 — that is, from a confident negative to a null. The structural prior demonstrably matters. What it did not do is produce per-gene skill, so a correct prior on one block is necessary but not sufficient; the intra-layer blocks remain pseudo-random draws, where a gene’s identity still enters the model only through its position. Four inputs are available at different cost. Reaction-level adjacency (KEGG, Reactome) or high-confidence interaction scores (STRING) would replace the regulatory and metabolic blocks directly. A curated compartment assignment would replace the present partition. Most valuable, and cheapest, is the stoichiometry: **S** currently encodes three moiety pools of which the rows without a metabolome member are filled arbitrarily, so real ATP/ADP and NAD^+^/NADH pools would make the conservation law biochemically true rather than only mathematically enforced.

Two cautions apply. First, enrichment statistics and curated adjacency are different objects. An enrichment score is computed from the same measurements the model is fitted to, so feeding it back as structure risks recovering an organisation that the analysis itself imposed; curated adjacency is independent of the data being fitted and is the safer input. Where enrichment is used it should follow the precedent already set here for the coherence term, whose weight is held at *λ*_coh_ = 0.01 precisely so that recovered structure remains a measured outcome. Second, real interaction networks are denser and far more hub-heavy than the surrogate, so a sparsification threshold becomes a methodological choice that must be reported rather than assumed.

The experiment this suggests is self-controlled and inexpensive: train the same architecture twice, once on the surrogate graph and once on curated adjacency, and compare per-gene skill. For the cross-layer block we have now run it. The surrogate template agreed with the true gene→protein correspondence of these pools on 12 of 66 edges, and replacing it changed the per-gene correlation as described above — a measurable effect on the metric, but not one that produced per-species skill. The same comparison on the intra-layer blocks remains to be done, and the surrogate is a degree-matched random control of exactly the kind it requires.

### C. Scale of a Whole-Cell Twin

It is useful to state what this model would become at whole-cell coverage, because the answer is not the one the O(*N* ^2^) figure of Section III H suggests. Taking a mammalian tri-omic inventory of *N* ≈ 4.4 × 10^4^ pools — roughly 2 × 10 transcripts, 2 × 10^4^ proteins, 3 × 10 metabolites and 10 lipid species — and holding the rhythmic fraction at the value the gate returns here (27%), the state is **x** ∈ R^79^ ^640^, and an interactome at realistic mean degree carries |*E*| ≈ 2.6 × 10^5^ edges.

At that size the representation choice dominates every other consideration. Materialised densely, **J** occupies 6.3 × 10^9^ entries, or 25 GB at single precision; stored on its biological support it occupies 5.3 × 10^5^ nonzeros, about 6 MB. The consequence for the trainable parameters is sharper still, and it falls on one component. The per-block heads that emit **J** are dense in the block dimension, predicting every upper-triangle entry, which is O(*N* ^2^) parameters: 1.2 × 10^11^ at whole-cell coverage, some 496 GB, comparable in size to a frontier language model. Predicting only the |*E*| entries that survive the mask instead gives 3.4 × 10^7^ parameters for the same operator, and 5.1 × 10^7^ for the whole model — roughly 200 MB, a single device. The reduction is a factor of 2.4 × 10^3^ and it costs nothing biologically, since the entries it declines to predict are masked to zero in any case. The energy network is unaffected by scale at all: its attention weights are shared across nodes, so it remains at 1.2 × 10^6^ parameters whether *N* is 10^2^ or 10^5^.

A whole-cell twin is therefore not compute-limited. Under the masked parameterisation a forward and backward pass costs of order 10*^−^*^1^ TFLOP at batch 48, with activation memory near 2 GB — the doubling relative to a conventional network being the price of retaining the gradient graph so that the loss can differentiate through ∇H.

The more interesting quantity is identifiability, and it runs opposite to intuition. Free parameters per observation, for a design of 24 timepoints, three replicates and ten conditions, is approximately 18 at the *N* = 100 scale of this study and approximately 1.6 at whole-cell coverage. The reason is that measurements scale with *N* — an assay reports every pool at once — while masked parameters scale with |*E*| ∝ *N*, and the fixed energy core is amortised over four hundred times as many nodes. *The configuration reported in this paper is the most over-parameterised point on that curve.* This reframes both null results: at eighteen parameters per observation, neither per-species generalisation nor identification of individual couplings should be expected, and the two nulls are evidence about this regime rather than necessarily about the architecture. The reframing is not a way of setting the nulls aside — it is a specific, falsifiable attribution, and it predicts that denser sampling should move both metrics while a larger network should not. Reaching parity at whole-cell coverage requires on the order of 38 timepoints, which is a demanding but ordinary circadian design.

We note one implication for a quantum realisation of the same construction. The classical bottleneck that motivates it is the *dense* O(*N* ^2^) representation; under the masked parameterisation above, the classical model remains tractable well beyond 10^5^ pools. The case for a quantum representation should therefore rest on the joint state and its correlations rather than on classical operator storage.

### D. What the Mixed Verdict Supports

The evidence reported here separates cleanly into claims of different strength, and conflating them would misrepresent the model. Structural properties hold by construction and require no data. System-level behaviour is supported: stability across ten seeds, held-out forecasting, and an aggregate cross-omic lag of 5.74 h against a measured 4.90 h from proteome timing the model was never fitted to. Two things are *not* supported. Per-species prediction: the per-gene lag correlation is indistinguishable from zero and its sign is unstable across seeds. And identification of network topology: recovery of withheld central-dogma edges sits at chance.

The two nulls share one likely cause, and it is a property of the data rather than of the formalism. Section V C shows the configuration used here is the most over-parameterised point on the scaling curve — of order 10^6^ parameters against 24 timepoints — so the dynamics are constrained enough to be stable and to extrapolate a trajectory, but not enough to localise which couplings produce them. That is an identifiability statement, and it predicts that denser temporal sampling, not a larger network, is what would move these two metrics. A reader shown the passivity and forecasting results without the two nulls would form a materially wrong impression of what this twin can do, which is why all five appear in Section IV and in Table II.

What distinguishes the framework is therefore not accuracy. It is that each of its claims can fail in a specified way: an exponent to be measured, an operator property to be violated, a lag to be mistimed, a conserved quantity to drift. A model whose output is a score can be inaccurate; it cannot be refuted. Developed against richer data, the realistic prospect is not a simulator that predicts every measurement but a theory of which interventions can and cannot work, and why — a narrower claim than a virtual cell, and a more defensible one.

## VI. CONCLUSION

We have presented a **composite, multi-clock, compartmental GNN-pHNN** for multi-omic cell dynamics. The framework rests on four elements:

*a. 1. Functional compartments with a composite Hamiltonian.* The storage function decomposes as H = ∑*_c_* H*_c_* + H_int_ over five functional compartments (core clock, redox, energy, signalling, biosynthesis), each a biological process rather than a measurement layer. This gives an energy accounting that maps onto how the cell is organised, and it lets passivity be certified compartment by compartment.

*b. 2. A clock bank of mechanistically distinct oscillators.* The model carries two genuinely distinct clocks — the transcription-dependent circadian TTFL and the transcription-independent redox rhythm — coupled through a zero-net-power NAD^+^/SIRT1 modulated port. A per-clock rhythmicity gate assigns every rhythmic pool to the correct clock from its spectrum alone (Figure 8), so multi-clock structure is inferred, not imposed.

*c. 3. Structure-preserving connectome and dissipation.* The interconnection is block-diagonal over compartments plus inter-compartment mass bonds and modulated ports, with the central dogma hard-wired and **R** = diag(*r_j_*) ≥ 0 initialised from measured degradation rates. Skew-symmetry and **R** ≥ 0 hold exactly. The learned connectome does *not*, however, recover withheld central-dogma edges on real mouse-liver data (AUROC 0.50 ± 0.13, nine scoreable seeds): the structure is imposed correctly, but 24 timepoints do not identify which edges carry it.

*d. 4. A falsifiable prediction with thermodynamic closure.* The composite loss enforces kinematic accuracy, global *and* per-compartment passivity, the power balance, conservation, and homeostasis. On real data the trained model satisfies max Ḣ = −1.7 ×10*^−^*^3^ ≤ 0 with zero violations across ten seeds. The turnover-sets-lag prediction Δ*φ* = arctan(*ω/k*_deg_), tested against author-measured transcript→protein lags with degradation rates fixed from published half-lives, reproduces the *aggregate* cross-omic lag (5.74 h predicted vs 4.90 h measured) while leaving the per-gene variation unresolved (*r* = 0.06 ± 0.27) — an honest partial confirmation that the loss was never told to satisfy, and whose limit (a layer-level degradation prior) is explicit.

Because these four elements are carried by the parameterisation rather than by the objective, they survive editing: a twin whose topology, dissipation, set-point or ports are deformed is still passive, still conservative, and still compartmentally decomposed. That is what makes this a *reference* twin rather than one fit among many, and it is the property that any account of specialisation or of disease built on this framework would rest on.

### A. Limitations and Future Directions

The storage function H is a Lyapunov function, not a directly measurable cellular energy; its absolute value carries no physical units. The results reported here are obtained on a real mouse-liver tri-omic dataset assembled from three separate public studies [24, 26] and Metabolomics Workbench, matched on circadian phase rather than same-animal sampling. This cross-cohort construction is the central caveat: the layers share a phase reference but not a biological sample, which limits any claim about coordinated same-cell dynamics. Two concrete data limits shaped the results above. First, degradation rates enter only as layer-level priors from published half-lives [28], so the cascade prediction is correct in aggregate but cannot resolve per-gene timing — closing this gap requires per-gene degradation measurements. Second, the public metabolome timecourse (six points over one 24 h cycle) is too sparse to resolve the rhythms the internal redox cascade needs, leaving that test inconclusive; a denser metabolome timecourse (≥ 8 points over ≥ 48 h) would resolve it.

A third limit is temporal resolution rather than layer coverage, and it bounds the two null results directly. Twenty-four timepoints against a parameter count of order 10^6^ place this study at the over-parameterised end of the identifiability curve of Section V C, and both the per-gene correlation and edge recovery are consistent with that regime rather than with a failure of the port-Hamiltonian constraint — which holds numerically throughout. We state the attribution as a prediction so that it can be refuted: denser temporal sampling should move both metrics, and a larger network at the same sampling should not. The edge-recovery null carries a further statistical caveat. The reference set contains 27 gene→protein edges, so a withheld fraction leaves at most a handful of positives; by the Hanley–McNeil variance a true AUROC below ≈ 0.79 is not separable from chance at this size. The null therefore excludes a large recovery effect and not a modest one, and a decisive test needs a denser reference network as well as an architectural change, since only the cross-layer block carries the additive deviation term through which a withheld edge can be scored at all.

Four directions follow, in order of priority:

1. **Same-cohort and denser multi-omics**: replace the cross-cohort assembly with matched-sample circadian multi-omics and a denser metabolome timecourse, enabling the internal dual cascade to be scored and the same-cell dynamics to be tested directly.
2. **Per-gene degradation rates**: fit or measure gene-specific *k*_deg_ so the cascade predic- tion can be tested per gene, not only in aggregate.
3. **A decisive topology-identification test**: score edge recovery against curated interaction networks (KEGG, Reactome, STRING) rather than a 27-edge correspondence, and add an additive deviation term to the intra-layer blocks so a withheld regulatory or metabolic edge can be scored. Both are required before the present null can be strengthened or overturned.
4. **Pharmacological in-silico screening and modular composition**: use the compartment-targeted ports to simulate drug action, and exploit the port-Hamiltonian compositionality guarantee to interconnect multiple cell instances through shared metabolic ports.

The composite compartmental GNN-pHNN establishes a rigorous, interpretable, and thermodynamically consistent foundation for a **Classical Digital Twin** of the living cell. What it does not yet establish is quantitative prediction at species resolution, and the results reported here locate that gap in the density of the available time courses rather than in the formalism — a claim we have stated in a form that further data can refute.

## Data Availability

All data used in this study are publicly available. The transcriptome is NCBI GEO accession GSE54650 [24, 25]; the proteome and the author-measured transcript→protein phase lags are from Robles, Cox & Mann 2014 [26] (PLoS Genetics 10(1):e1004047 and its Table S4); the metabolome is NIH Metabolomics Workbench accession ST002079 [27] (doi:10.21228/M87125). Protein and mRNA degradation half-lives used to fix *k*_deg_ are from Schwanhäusser et al. 2011 [28]. The derived, harmonised tri-omic bundle is assembled from the public datasets cited above, from which the full retrieval provenance for each layer can be traced. The code used in this study, including the model, the training pipeline and the scripts that regenerate every figure and reported number, is available at https://github.com/quantum-omics/Classical-Virtual-Omics.

